# A non-canonical function of Dicer fuels inflammation driven by self-dsRNAs

**DOI:** 10.64898/2026.10.29.761885

**Authors:** Laurens R. ter Haar, Sabine de Lange, Dennis Gravekamp, Annick C. Kooij, Jasper W. de Wolf, Yanniek Asschert, Timo Oosenbrug, Maarten W. Paul, Peter A. van Veelen, Willeke M.C. van Roon-Mom, Hailiang Mei, Annemarthe G. van der Veen

## Abstract

Self-derived double-stranded RNA (self-dsRNA) acts as potent triggers of autoinflammation and damage-associated molecular patterns in cancer and infection. However, the upstream processes controlling self-dsRNA homeostasis remain unclear. We uncover a non-canonical role for human Dicer in driving cell-intrinsic immunity to self-dsRNAs that arise from editing or splicing defects, epigenetic deregulation or mitochondrial stress. Dicer loss abolishes type-I interferon induction, PKR activation, and OAS-RNase L activation in response to self – but not exogenous – dsRNA in multiple cell types, including a clinically-relevant hiPSC model of neuroinflammation. Dicer interacts with G3BP1 and LGP2 to regulate self-dsRNA availability and recognition, respectively, independently of miRNAs. As immunostimulatory self-dsRNAs fail to accumulate and activate dsRNA sensors in absence of Dicer, these findings position Dicer as a central regulator of self-dsRNA sensing.

**Summary figure:** 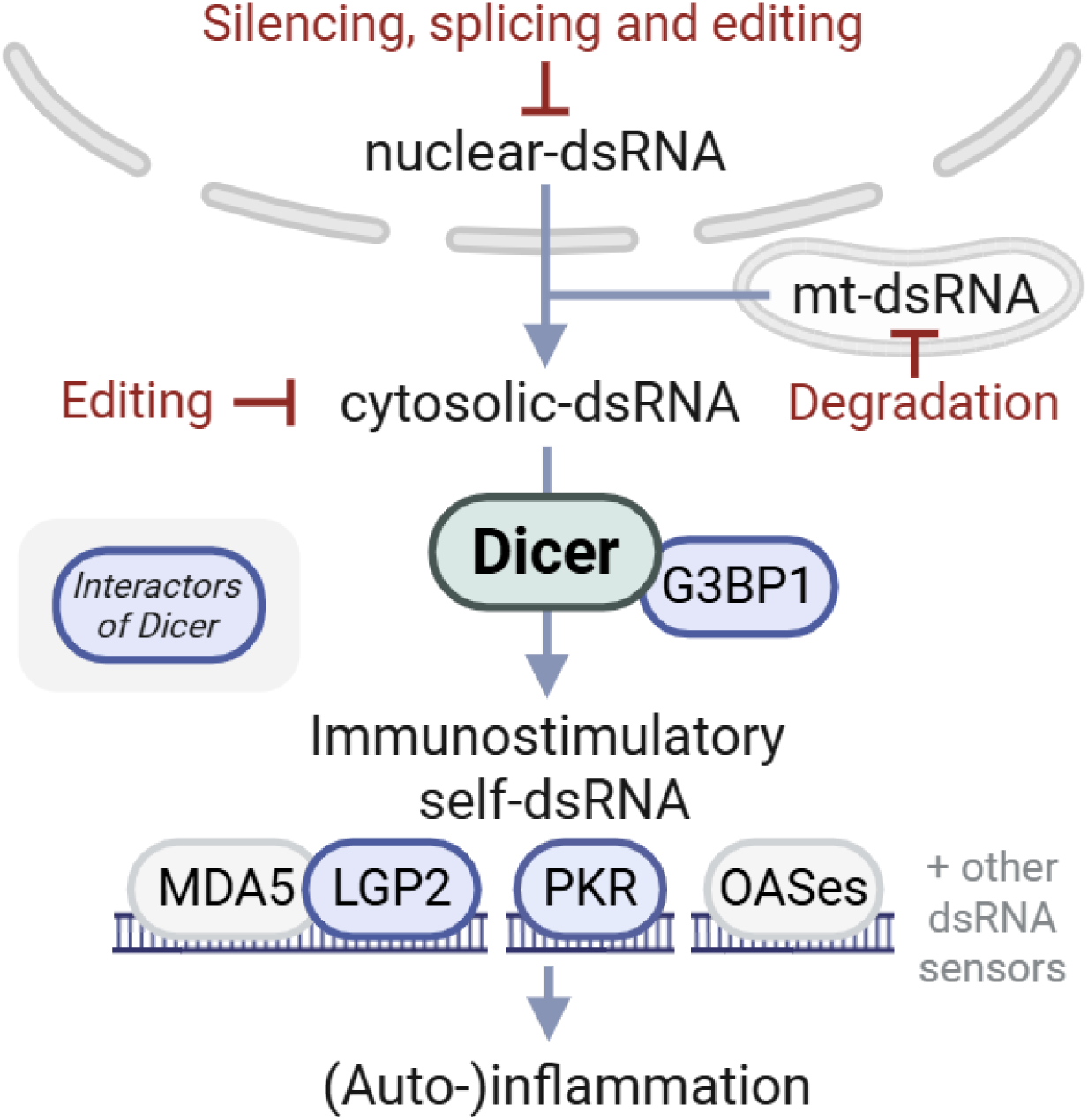

Self-derived dsRNAs are potent triggers of innate immune activation that accumulate due to cellular dysfunction, such as mitochondrial stress, epigenetic deregulation or editing and splicing defects. Dicer and G3BP1 are jointly required for the cytosolic accumulation of self-dsRNA, and the activation of downstream innate immune dsRNA sensors, including Dicer interactors LGP2 and PKR. Self-dsRNA driven inflammation occurs in diverse pathological contexts – including neurodegeneration, autoimmunity, and cancer – thus making Dicer and its interactors promising targets for therapeutic intervention.

Created in BioRender. Ter haar, L. (2025) https://BioRender.com/40ztdl0

## Main text

Cell-intrinsic innate immune pathways form a barrier against viral infection. Double-stranded RNA (dsRNA) is abundantly present in viral particles or generated during the replication cycle of RNA viruses. Cells have evolved cytosolic sensors that are activated by viral dsRNA. Upon activation, dsRNA sensors induce the dsRNA response, that includes the type I interferon (IFN-I) response, the integrated stress response, inhibition of cell growth, and cell death.(*1, 2*) While traditionally viewed as a hallmark of viral infection, dsRNA can also arise from endogenous cellular processes such as bidirectional transcription or expression of repetitive retroelements (REs) which can form intramolecular base-paired or bona fide intermolecular dsRNA (collectively referred to as self-dsRNA).(*1, 3–5*) The accumulation of self-dsRNAs can aberrantly activate dsRNA sensors and cause autoinflammation and immunopathology.(*1, 2, 4–6*) To circumvent this, cells use counteractive measures such as modification, degradation, or epigenetic repression of immunostimulatory self-dsRNA.(*3, 6*)

Failure to suppress self-dsRNA-driven immune activation is a major disease-causing mechanism. This is best exemplified by patients with inborn errors of immunity known as type I interferonopathies, which includes the prototypical disorder Aicardi-Goutières Syndrome (AGS).(*2, 7, 8*) AGS patients develop severe neuroinflammatory symptoms and skin manifestations due to single gene defects. These include loss-of-function mutations in the RNA editing enzyme adenosine acting on RNA (*ADAR*), which lead to an accumulation of unedited self-dsRNAs that drive a pathogenic dsRNA response. (*2, 7–9*)

Immunostimulatory self-dsRNA can derive from mitochondrial or nuclear sources. Bidirectional transcription in mitochondria generates large amounts of dsRNA that is degraded by the RNA degradosome complex containing the ribonuclease polynucleotide phosphorylase (PNPase; encoded by *PNPT1*).(*6, 10*) Under certain circumstances, mitochondrial-derived dsRNA (mt-dsRNA) can accumulate, escape to the cytoplasm and drive a dsRNA response. Cytosolic mt-dsRNA is associated with osteoarthritis, renal injury, Huntington’s disease, and cellular senescence.(*11–14*) Hypomorphic mutations in *PNPT1* lead to a constitutive IFN-I response and a type I interferonopathy.(*8, 10*)

Defective splicing may also increase the intracellular pool of self-dsRNA. Heterogeneous nuclear ribonucleoprotein C (hnRNPC) is a ubiquitously expressed nuclear protein that controls alternative splicing.(*15*) Loss of hnRNPC causes an accumulation of self-dsRNAs originating from pre-mRNA introns containing Alu-repeat sequences, which leak into the cytosol and activate an IFN-I response.(*16, 17*) Finally, epigenetic derepression of REs causes self-dsRNAs to accumulate (*18–20*). For example, inactivation of the human silencing hub (HUSH) complex through depletion of M-phase phosphoprotein 8 (MPP8, encoded by *MPHOSPH8*) leads to upregulation of certain REs, which drive a spontaneous IFN-I response.(*21, 22*)

DsRNA of self or viral origin activates cytosolic dsRNA sensors, including the RIG-I-like-receptors (RLRs) RIG-I (retinoic acid–inducible gene I), MDA5 (melanoma differentiation factor 5), and LGP2 (laboratory of genetics and physiology 2). RIG-I recognizes short blunt-end dsRNA containing a 5’di-or triphosphate moiety, while MDA5 and LGP2 cooperatively sense long dsRNA onto which they oligomerize to form filaments.(*6, 23*) Self-dsRNAs that accumulate in the absence of hnRNPC or MPP8 are reported to activate both MDA5 and RIG-I, while loss of PNPase or ADAR1 activates MDA5.(*5, 10, 16, 17, 22, 24–28*) LGP2 is essential for MDA5-mediated IFN-I induction in ADAR1 deficiency, but its role in other contexts of self-dsRNA accumulation remains unexplored.(*5, 28*) RLR activation leads to phosphorylation of TBK1, and phosphorylation and nuclear translocation of the transcription factors IRF3 and NF-kB, which drive transcription of type I or type III IFNs and pro-inflammatory cytokines, respectively. Secreted IFNs activate the IFN receptor (IFNAR) in an autocrine and paracrine manner, followed by JAK-STAT signalling and the upregulation of hundreds of ISGs, encoding proteins with antiviral activity.(*2, 23*)

Besides RLRs, cytosolic dsRNA can activate additional dsRNA sensors, including PKR, ZBP1 and OAS1-3. Dimerization of PKR onto dsRNA leads to PKR-mediated autophosphorylation and phosphorylation of eIF2a, inducing the integrated stress response, the nucleation of stress granules (SGs) and inhibition of cell growth. ZBP1 recognizes dsRNA in its unusual left-handed Z-conformation, triggering NF-kB activation and cell death. OAS1–3 detect dsRNA and activate the ribonuclease RNase L, which degrades both viral and cellular RNA, ultimately inducing cell death. The expression of these dsRNA sensors is upregulated by type I interferon (IFN-I) signaling. Hence, even low basal activation of RLRs can initiate a positive-feedback loop that amplifies the expression of dsRNA sensors, reinforcing their involvement in disease-associated immune responses.(*2, 29*)

While many studies have elucidated how the cytosolic dsRNA sensors RLRs, PKR, ZBP1, and OAS1-3 detect self-dsRNA (*1, 2, 4–6, 30*), the upstream mechanisms governing self-dsRNA formation and accumulation prior to detection remain unclear. Insight into these mechanisms may reveal attractive strategies for therapeutic targeting to specifically limit pathological responses to self-dsRNA, without compromising antiviral immunity. We set out to study the role of the type III endoribonuclease Dicer, another cytosolic dsRNA-binding protein, in the context of autoinflammation and interferonopathies. Dicer is evolutionarily related to the RLRs and shares a conserved N-terminal helicase domain. In plants and invertebrates, Dicer acts as a key antiviral effector by mediating RNA interference (RNAi).(*31–33*) In mammals, however, Dicer exhibits limited antiviral RNAi activity in IFN-competent, differentiated cells and instead plays a central role in gene regulation via the microRNA (miRNA) pathway.(*31–34*) This functional shift has been attributed to Dicer’s helicase domain, which restricts its processing of long dsRNA and favors the cleavage of imperfect hairpin structures such as pre-miRNAs.(*35–37*) Moreover, the IFN-I pathway is functionally incompatible with the antiviral RNAi pathway.(*34, 38–41*) Over the course of evolution, the RLR-dependent IFN-I pathway may have supplanted the antiviral role of Dicer. This transition could have necessitated a reduction in Dicer’s ability to degrade long dsRNA, thereby preserving these molecules as substrates for cytosolic dsRNA sensors. Of note, previous studies revealed that human Dicer binds to RE-element-containing RNA and mt-RNA, the proposed sources of self-dsRNAs.(*42*) Nevertheless, whether and how Dicer precisely impacts on the activation of dsRNA sensors by self-dsRNA remains unanswered.

We uncover that Dicer acts as a central regulator of self-dsRNA homeostasis and innate immune activation by self-dsRNAs in a miRNA-independent manner. Dicer interacts with G3BP1 to regulate immunostimulatory self-dsRNA availability, and subsequently facilitates recognition of self-dsRNAs via its association with LGP2 and PKR. These insights reveal an unexpected aspect of innate immune activation by self-dsRNAs with important implications for novel therapeutic strategies.

## Results

### Dicer is required for the activation of MDA5 and LGP2 upon ADAR1 ablation

To investigate a role for the endoribonuclease Dicer in the context of self-dsRNA sensing, we depleted ADAR1 using siRNAs in human embryonic kidney 293 cells (HEK293) to induce a sterile, MDA5/LGP2-dependent IFN-I response. As HEK293 cells are naturally devoid of functional LGP2 protein levels, we used cells that were stably transduced with a lentivirus encoding human LGP2 (HEK293-L).(*5*) Loss of ADAR1 induced robust upregulation of transcripts encoding IFN-β (IFN-I), IFIT1 and ISG15 (two prototypical ISGs), measured by RT-qPCR (Fig. 1A and Fig. S1A-B). Remarkably, Dicer co-depletion using independent siRNAs completely abolished this robust IFN-I response (Fig. 1A and Fig. Fig. S1A-B).

**Figure 1:**
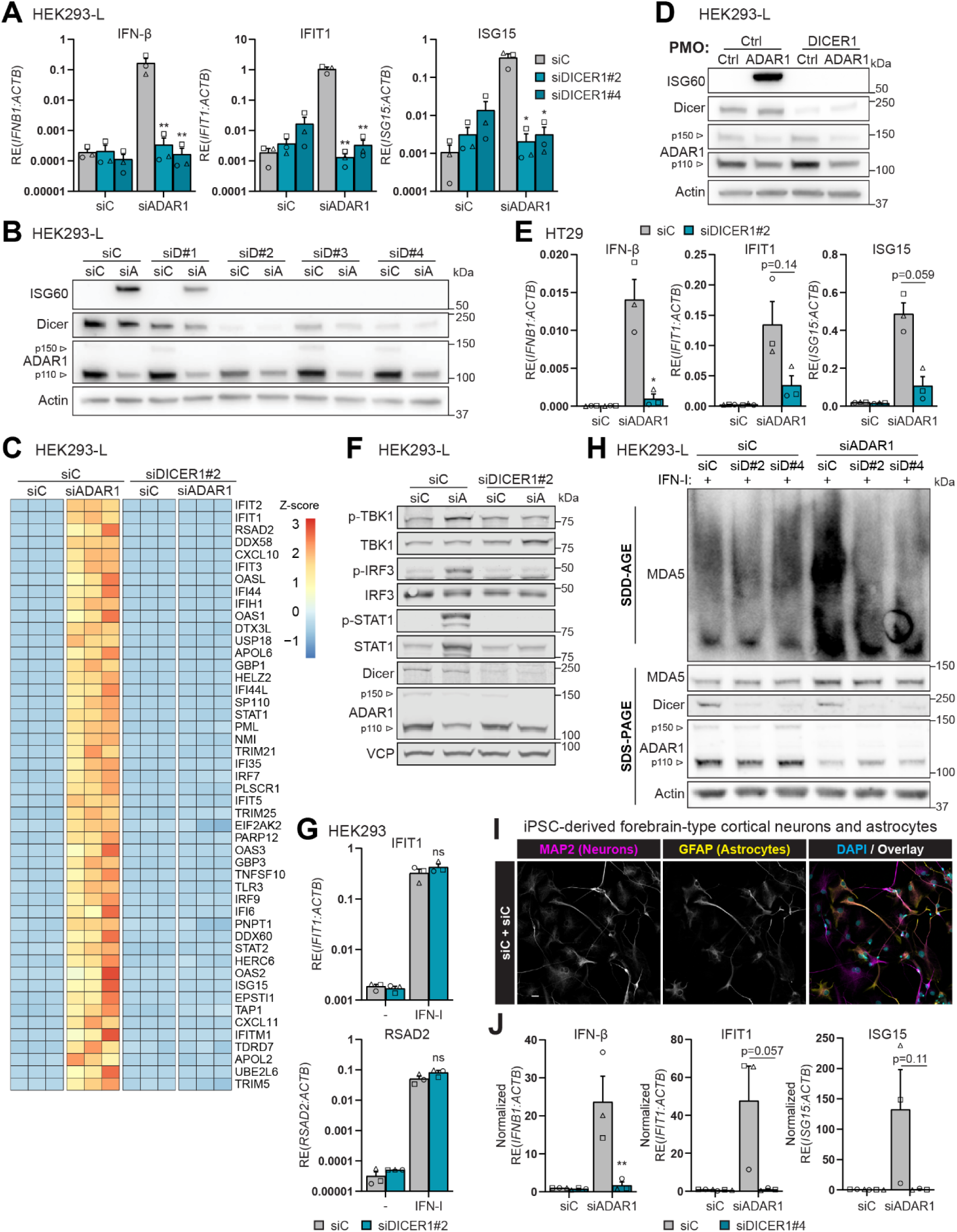
Dicer is required for the MDA5/LGP2-driven IFN-I response following ADAR1 ablation. **(A-C)** HEK293-L cells 72h post-siRNA transfection (n=3). **(A)** IFN-I response by RT-qPCR. **(B)** ISG protein levels by immunoblot. **(C)** Heatmap of top-50 ISG genes by RNA-sequencing. **(D)** HEK293-L cells 72h post-PMO treatment analysed by immunoblot (n=3). **(E)** HT29 cells 72h post-siRNA transfection, IFN-I response assessed by RT-qPCR (n=3). **(F)** HEK293-L cells 72h post-siRNA transfection analysed by phospho-immunoblot (n=4). **(G)** HEK293 cells pretreated with siDICER1#2 for 24h, then stimulated for 16h with recombinant IFN-I and assessed by RT-qPCR (n=3). **(H)** HEK293-L cells 72h post-siRNA transfection, subjected to SDD-AGE to determine MDA5 oligomerization or SDS-PAGE for protein levels. Cells were treated with recombinant IFN-I 24h before harvest to equalize MDA5 levels (n=4). **(I-J)** Healthy hiPSC-derived neural progenitor cells differentiated for 7 days before 15 days of maturation into forebrain-type cortical neurons and astrocytes. Cells were siRNA treated 5 days before harvest (n=3). **(I)** Differentiation verified by immunofluorescence; scale bar: 20 µm. **(J)** IFN-I response by RT-qPCR. Data shown as means ±SEM (RT-qPCRs) or representative images/blots. Statistical tests: one-way ANOVA (A), paired t-test (E, G and J). ns, not significant; \**P*<0.05; **P<0.01. siA, siADAR1; siD, siDICER1; siC, siControl.

Similarly, immunoblotting of ADAR1-depleted cells revealed attenuation of ISG60 protein expression upon co-depletion of Dicer using four individual siRNAs (Fig. 1B). The extent of IFN-I reduction correlated with Dicer knockdown efficiency (Fig. 1B and S1A-B). RNA sequencing showed that depletion of Dicer globally abolished ISG induction after ADAR1 loss (Fig. 1C and S1C). Note that the loss of Dicer did not impact on the knockdown efficiency of ADAR1, since 21-mer siRNAs are directly loaded into the RNA-induced silencing complex (RISC) and bypass the need for Dicer processing, unlike miRNAs or longer Dicer-substrate siRNAs.(*43, 44*) We recapitulated these findings with an independent method by depleting Dicer and ADAR1 using phosphorodiamidate morpholino oligomers (PMOs, that induce exon skipping to interfere with correct translation) delivered via Endo-Porter or cell-penetrating vivo-PMOs (vPMOs) (Fig. 1D and S1D-E). Consistently, loss of Dicer inhibited the IFN-I response upon ADAR1 depletion in the colorectal cancer cell line HT29, which expresses functional levels of endogenous LGP2 (Fig. 1E and S1F).(*5*)

Analysis of downstream signalling events revealed that phosphorylation of TBK1, IRF3, and STAT1 upon loss of ADAR1 were all abolished by simultaneous Dicer depletion in HEK293-L cells (Fig. 1F). Similarly, loss of Dicer attenuated phosphorylation of TBK1 and IRF3 in ADAR1-deficient LGP2-expressing A549 lung epithelial cells (A549-L) (Fig. S1G). Since MDA5 and LGP2 are ISGs, we asked whether Dicer may indirectly affect their activation through regulating IFNAR signalling and the downstream induction of ISGs.

However, Dicer depletion did not impact on ISG induction in HEK293 cells treated with recombinant IFN-I (Fig. 1G and S1H). Furthermore, IFN-I treatment of ADAR1-depleted HEK293-L and A549-L cells did not restore phosphorylation of TBK1 and IRF3 after Dicer depletion (Fig. S1I-J). MDA5 activation occurs through oligomerization along dsRNA ligands.(*45*) To study the impact of Dicer on MDA5 oligomerization directly, we visualized MDA5 aggregates by semi-denaturing detergent agarose gel electrophoresis (SDD-AGE).(*46, 47*) Consistently, MDA5 oligomerization in ADAR1-depleted HEK293-L cells is abolished by Dicer co-depletion (despite using recombinant IFN-I to boost and equalize endogenous MDA5 expression levels) (Fig. 1H). Together, these data demonstrate that Dicer is required for MDA5/LGP2-dependent sensing of self-dsRNA following ADAR1 ablation.

Patients with the prototypical type I interferonopathy AGS develop severe neuroinflammatory symptoms. We modelled neuroinflammation by ablating ADAR1 expression with siRNAs in mixed human induced pluripotent stem cells (hiPSC)-derived forebrain-type cortical neurons and astrocytes differentiated from healthy human neural progenitor cells (Fig. 1H). The strong IFN-I response induced after ADAR1 ablation was completely abrogated by Dicer co-depletion, implicating Dicer as a crucial factor in the neuropathological IFN-I response of AGS (Fig. 1I and S1K).

### Dicer is not required for the IFN-I response to exogenous dsRNA ligands

We subsequently examined whether Dicer is involved in IFN-I induction in response to a panel of exogenous RLR ligands. However, activation of RLRs with ligands that are predominantly sensed by RIG-I (3p-hpRNA or 3p-IVT dsRNA) or MDA5 (HMW poly(I:C)) induced an IFN-I response in HEK293 cells in a Dicer-independent manner (Fig. 2A and S2A), consistent with literature.(*40*) To avoid the potential confounding effects of Dicer’s role in antiviral RNAi, we did not investigate IFN-I induction during the course of a bona fide viral infection. However, in line with the above findings, Dicer did not impact on the MDA5/LGP2-dependent IFN-I response following transfection of RNA containing encephalomyocarditis virus (EMCV) genomes and replication intermediates into HEK293-L cells (Fig. 2B and S2B).(*5*) We conclude that Dicer is not involved in IFN-I induction upon activation of dsRNA sensors by exogenous ligands.

**Figure 2:**
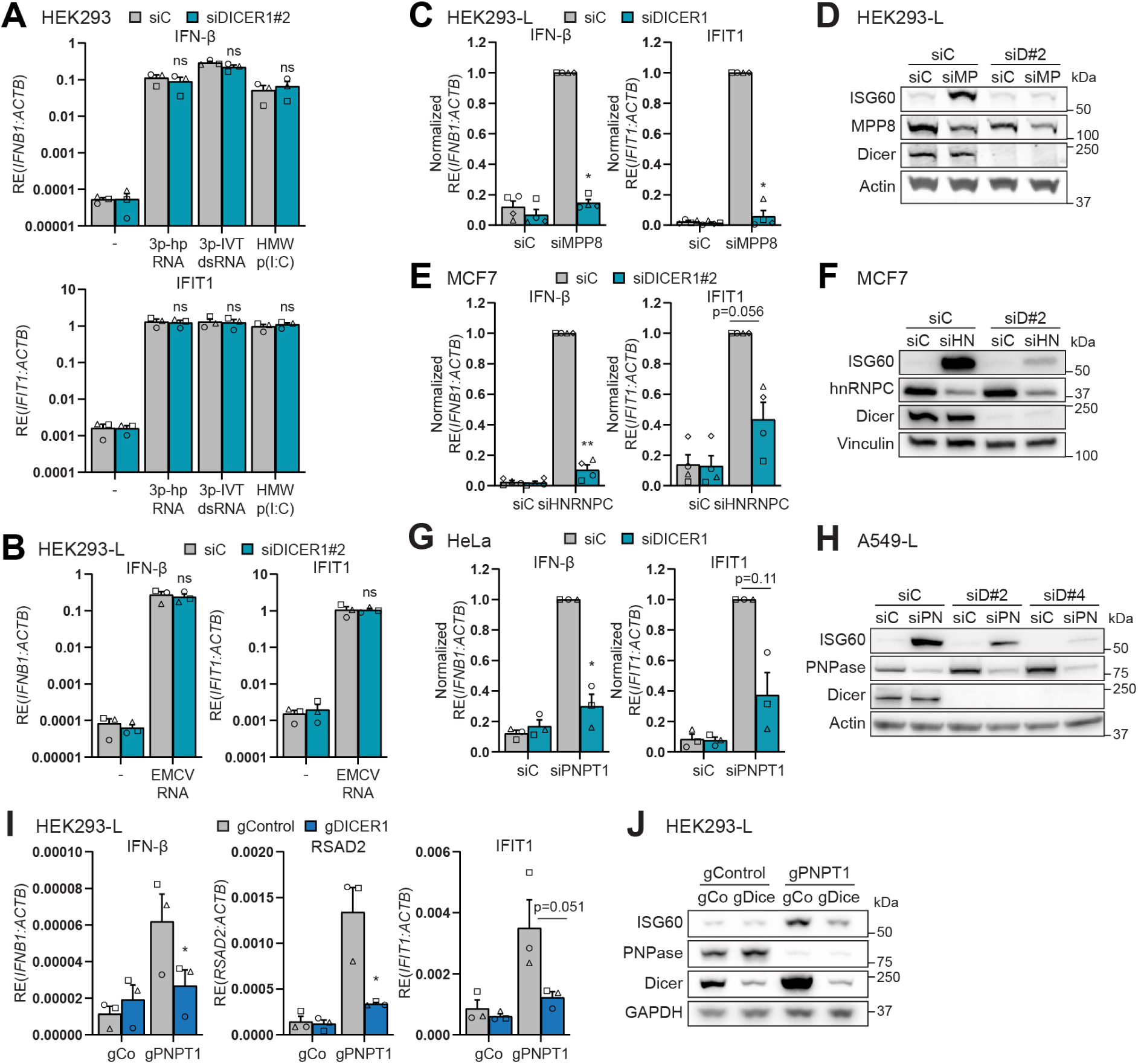
Dicer is required for cell-intrinsic immunity to self-dsRNA of both nuclear and mitochondrial origin. **(A-B)** Cells pretreated with siDICER1#2 for 24h, then transfected for 16h with synthetic RLR ligands in **(A)**, or RNA from EMCV-infected cells (Dipyridamole was used to supress viral replication) in **(B)**. IFN-I response assessed by RT-qPCR (n=3). **(C-H)** Cells transfected with siRNAs targeting MPP8 for 6 days **(C-D)**, hnRNPC for 3 days **(E-F)**, or PNPase for 4 days **(G-H)**. IFN-I response monitored by RT-qPCR, transcripts normalized to ACTB and relative to siC/siMPP8 (**C**; n=3), siC/siHNRNPC (**E**; n=4) or siC/siPNPT1 (**G**; n=3). Protein expression analysed by immunoblot (**D**; n=2, **F, H**; n=3). **(I-J)** HEK293-L cells 6 days post-nucleofection with Cas9 and gRNAs targeting *PNPT1* and/or *DICER1* (n=3). **(I)** IFN-I response by RT-qPCR. **(J)** Protein levels by immunoblot. gCo, gControl; gPN, gPNPT1. Data shown as means ±SEM (RT-qPCRs) or representative images/blots. Statistical tests: one-way ANOVA (A), paired t-test (B, C, E, G, I). ns, not significant; \**P*<0.05; **P<0.01. siD, siDICER1; siMP, siMPP8; siHN, siHNRNPC; siPN, siPNPT1; siC, siControl.

### Dicer is required for cell-intrinsic immunity to self-dsRNA of both nuclear and mitochondrial origin

To examine whether Dicer is broadly implicated in self-dsRNA sensing or whether its role is restricted to ADAR1 ablation, we employed several cellular models of autoinflammation driven by distinct sources of self-dsRNA. Ablation of MPP8 leads to the epigenetic derepression of RE expression and induction of the IFN-I response, mediated predominantly via MDA5 and partly RIG-I.(*22*) Notably, the absence of Dicer prevented the induction of an IFN-I response in HEK293-L cells upon MPP8 depletion (Fig. 2C-D and S2C). Loss of the splicing factor hnRNPC leads to an accumulation of Alu-containing introns that form self-dsRNAs and trigger an IFN-I response via MDA5 and RIG-I. (*16, 17*) We recapitulated these observations and demonstrated that the simultaneous depletion of Dicer attenuated the IFN-I response in hnRNPC-depleted MCF7 breast cancer cells (Fig. 2E-F and S2D). Finally, deficiency in mitochondrial PNPase activity causes an accumulation of mt-dsRNAs in the cytoplasm, which drive an MDA5-dependent IFN-I response. (*10, 11*) Reduction in mitochondrial PNPase activity by siRNA-mediated depletion of PNPase yielded a sterile IFN-I response that is dependent on the presence of Dicer in HeLa, A549-L and HEK293-L cells (Fig. 2G-H and S2E-H). These results were replicated in HEK293-L cells using recombinant Cas9 and gRNAs directed against *PNPT1* and *DICER1* to simultaneously knockout both genes (Fig. 2I-J and S2I). Cells were harvested six days later to avoid the detrimental consequences of chronic *DICER1* deficiency (including mild induction of ISGs that we first observe after 14 days of Dicer depletion, consistent with literature) (Fig. S2J-K).(*48, 49*) Knockout of *PNPT1* induced an IFN-I response, which was prevented by the simultaneous loss of *DICER1*. Altogether, these observations suggest that Dicer is broadly required for cell-intrinsic immunity to self-dsRNA of both nuclear and mitochondrial origin.

### Dicer is required for PKR and OAS1-3/RNase L pathway activation upon ADAR1 ablation

We next examined whether other cytosolic dsRNA sensing pathways also require the presence of Dicer. Self-dsRNAs that accumulate upon ADAR1 ablation are also sensed by PKR, leading to its autophosphorylation and eIF2a phosphorylation.(*24, 27, 28*) Furthermore, sensing of self-dsRNA by PKR, MDA5 and ZBP1 drives activation of inflammatory gene transcription factor NF-kB through degradation of its inhibitor IκBα.(*1*) Strikingly, Dicer depletion abrogated PKR autophosphorylation, eIF2a phosphorylation, and IκBα degradation following ADAR1 ablation in A549-L and HEK293-L cells (Fig. 3A and S3A).

**Figure 3:**
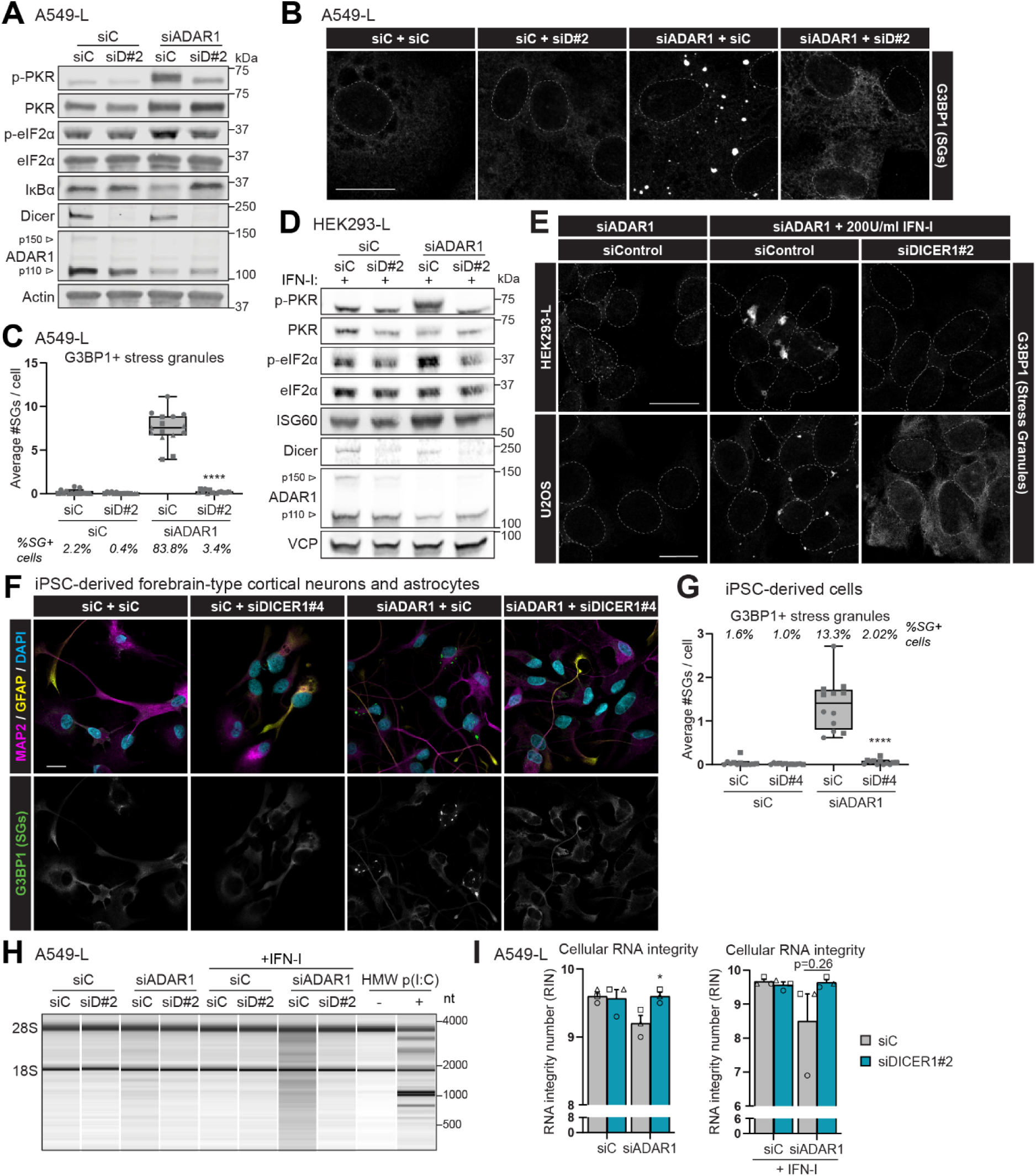
Dicer is broadly required for cell-intrinsic immunity to self-dsRNA following ADAR1 ablation. **(A)** A549-L cells 72h post-siRNA transfection analysed by phospho-immunoblot (n=3). **(B and E)** Stress granule formation in A549-L (**B**, n=6), HEK293-L (**E**, n=4) and U2OS (**E**, n=3) cells 72h post-siRNA transfection and/or 48h post-IFN-I treatment, visualized by G3BP1 immunofluorescence. Nuclear DAPI stain outlined. **(C)** Quantification of immunofluorescence in (B). 1624-2615 cells were quantified across 15 images per condition (n=3). **(D)** HEK293-L cells 72h post-siRNA and IFN-I treatment to upregulate PKR, analysed by phospho-immunoblot (n=4). **(F-G)** Healthy hiPSC-derived forebrain-type cortical neurons and astrocytes (differentiated as in Fig. 1I) harvested 5 days post-siRNA transfection. **(F)** Stress granule formation visualized by immunofluorescence (n=3). **(G)** Quantified across 12 images per condition (n=2). **(H-I)** A549-L cells 72h post-siRNA and/or 48h post-IFN-I or 6h post-HMW poly(I:C) transfection. **(H)** OAS1-3/RNase L activation measured by integrity analysis of total RNA extracts using Bioanalyzer electrophoresis (n=3). **(I)** Cellular RNA integrity quantified using RNA integrity number (RIN). Data shown as means ±SEM (RT-qPCRs and RIN values) or representative images/blots. Box plots showing interquartile range (IQR), median and whiskers (1.5×IQR), shapes represent biological repeats. Scale bars: 20 µm. Statistical test: paired t-test (C, G and I). \*\*\*\**P*<0.0001. siD, siDICER1; siC, siControl.

Phosphorylation of eIF2a leads to translation shutdown, driving SG formation through phase-condensation of SG nucleators on translationally stalled mRNAs.(*50*) Consistent with earlier reports, immunofluorescence of SG-nucleator G3BP1 revealed SG induction after ADAR1 depletion in A549-L cells (Fig. 3B-C).(*51, 52*) Simultaneous depletion of Dicer fully abrogated SG formation. We confirmed these were bona fide SGs – and not G3BP1^+^ RNase L-induced bodies – by staining for the SG-exclusive marker TIA1, which completely overlapped with G3BP1 staining (Fig. S3B).(*50*) To rule out the possibility that Dicer is indirectly affecting PKR activation through the loss of its upregulation as an ISG following MDA5/LGP2 activation, we treated ADAR1-depleted cells with exogenous IFN-I. IFN-I treatment did not restore phosphorylation of PKR and eIF2a after Dicer co-depletion in HEK293-L and A549-L cells (Fig. 3D and S3C). Dicer depletion also abolished SG induction in IFN-I-treated, ADAR1-deficient A549-L, HEK293-L and U2OS cells (Fig. 3E and S3D). Finally, we examined PKR activation in our hiPSC-derived model of AGS neuroinflammation. Consistently, PKR-induced SG assembly upon ADAR1 depletion was completely reverted by Dicer co-depletion (Fig. 3F-G). These data show Dicer is directly required for the activation of PKR in ADAR1-deficient cells.

Self-dsRNA accumulation upon ADAR1 depletion also activates dsRNA sensors OAS1-3, which trigger RNase L to degrade cellular RNAs.(*53, 54*) RNA integrity measurements revealed that RNase L was modestly activated upon loss of ADAR1 in A549-L cells (but not HEK293-L or HT29 cells), which was boosted by IFN-I treatment to upregulate OAS1-3 (Fig. 3H-I and S3E). Notably, Dicer co-depletion completely halted RNase L activation in ADAR1-depleted A549-L cells, irrespective of IFN-I treatment, showing that Dicer is also required for OAS1-3 activation in ADAR1-deficient cells. Collectively, these findings show that Dicer is broadly required for cell-intrinsic immunity to self-dsRNA following ADAR1 ablation.

### Dicer does not impact on self-dsRNA sensing through the miRNA pathway

We next investigated whether Dicer impacts on self-dsRNA sensing through its canonical role in miRNA biogenesis. The microprocessor complex consisting of Drosha and DGCR8 processes primary miRNAs into precursor miRNAs, which Dicer further cleaves into mature miRNAs.(*31*) Unlike Dicer, depletion of Drosha or DGCR8 in ADAR1-deficient HEK293-L cells, or Drosha depletion in ADAR1-deficient HT29 cells, did not affect the IFN-I response (Fig. 4A–B and S4A). Dicer processing generates mature miRNAs, which are loaded into the RISC to mediate translation inhibition or degradation of complementary target RNAs, primarily via Ago2, the only human Argonaute with endonucleolytic activity.(*31*) We knocked down Ago2 and ADAR1 using vPMOs (since Ago2 is required to establish siRNA-mediated knockdowns) in HEK293-L cells. Knockdown of Ago2 did not affect the IFN-I response in ADAR1-depleted cells (Fig. 4C-D and S4B). These data show that Dicer does not impact on the IFN-I response in ADAR1-deficient cells via the miRNA pathway.

**Figure 4:**
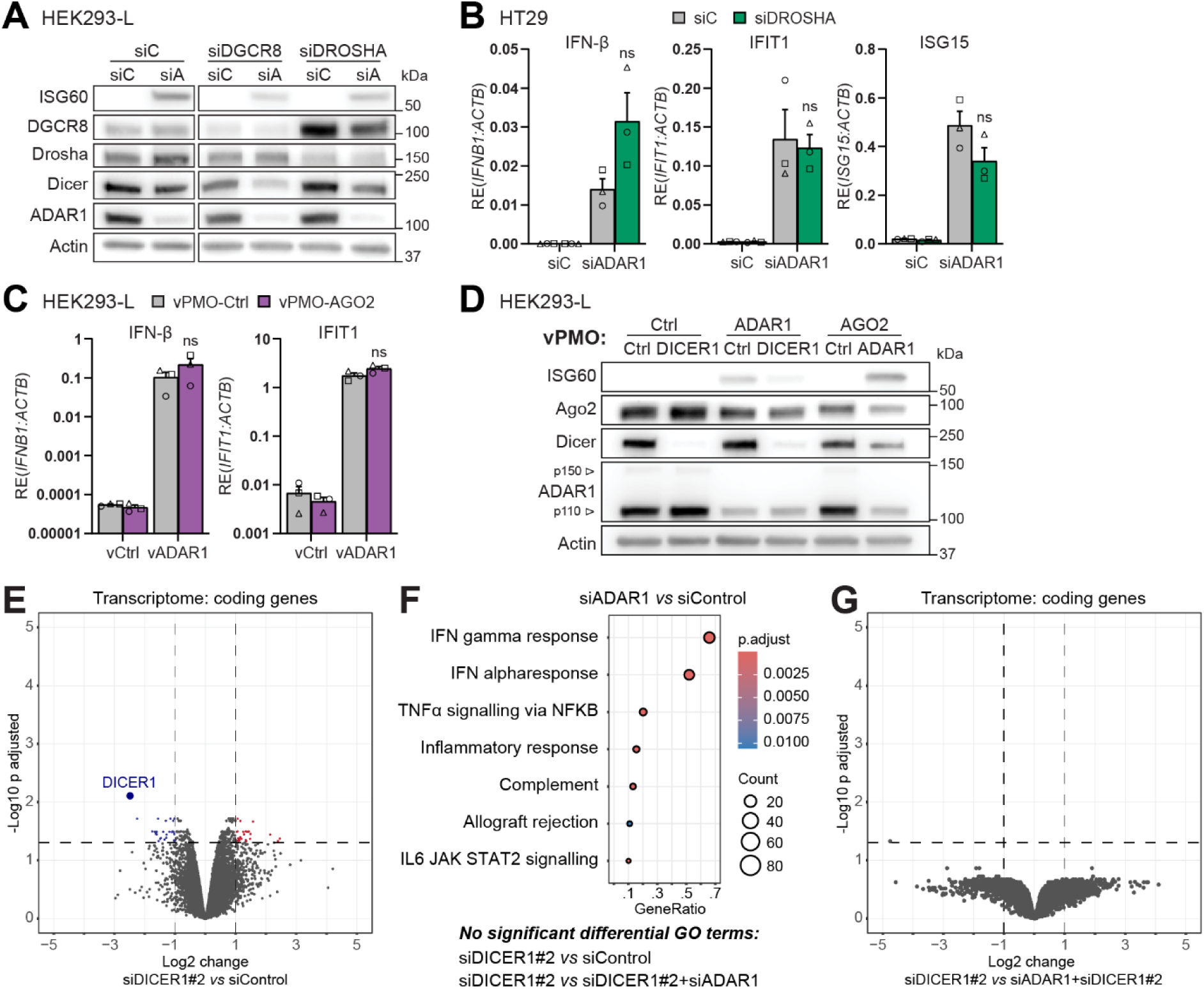
Dicer does not impact on self-dsRNA sensing through miRNA biogenesis or transcriptional changes. **(A)** HEK293-L cells 72h post-siRNA transfection analysed by immunoblot (n=3). **(B)** HT29 cells 72h post-siRNA transfection analysed by RT-qPCR (n=3). **(C-D)** HEK293-L cells 72h post-treatment with vPMOs (n=3). **(C)** IFN-I response by RT-qPCR. **(D)** Protein levels by immunoblot. **(E-G)** Transcriptomics of HEK293-L cells shown in Fig. 1C, 72h post-siRNA transfection (n=3). **(E and G)** Differential expression analyses. **(F)** GO-enrichment analyses. Data shown as means ±SEM (RT-qPCRs) or representative images/blots. Statistical tests: paired t-test (B-C). ns, not significant.

To further investigate whether Dicer impacts on self-dsRNA sensing at a transcriptional level, we examined our RNA sequencing dataset of ADAR1-and/or Dicer-depleted HEK293-L cells (Fig. 1C). Dicer depletion alone did not induce substantial changes in mRNA transcript levels during the investigated time frame (72 hrs) (Fig. 4E). Consistently, Gene Ontology (GO) enrichment analysis revealed no significantly affected pathways after Dicer depletion, unlike ADAR1-depleted cells, which revealed the expected IFN-I and pro-inflammatory signature (Fig. 4F). Furthermore, ADAR1/Dicer co-depletion caused no significant transcriptional changes on top of Dicer depletion alone (Fig. 4F-G), despite ADAR1’s role in modulating Dicer’s siRNA and miRNA processing activity.(*55–57*) Finally, aside from inflammatory genes and ISGs, which are no longer upregulated in ADAR1/Dicer co-depleted cells, ADAR1/Dicer co-depletion caused limited additional changes in mRNA expression compared to ADAR1-depleted cells (Fig. S4C). The minor impact of Dicer and ADAR1 depletion on global gene expression is not unexpected and likely reflects the partial and short-term depletion achieved through siRNAs. Altogether, these data suggest that Dicer does not impact on self-dsRNA sensing through its role in regulating gene expression via miRNA maturation.

### Self-dsRNA sensing does not remodel the global Dicer interactome

To investigate the mechanism underlying Dicer’s involvement in self-dsRNA sensing, we used proteomics to determine how the Dicer interactome is reshaped during self-dsRNA sensing. We immunoprecipitated FLAG-tagged Dicer from control and ADAR1-depleted HEK293-L cells, followed by mass spectrometry analysis (Fig. S5A). To discriminate between interactors that are due to bona fide alterations in protein-protein interactions versus those that reflect increased abundance due to their IFN-I inducibility, we also included ADAR1-depleted cells that were treated with the JAK-kinase inhibitor ruxolitinib to block the upregulation of inflammatory genes and ISGs.(*58*) We detected the well-known Dicer interactors AGO1-3, TRBP2 and PACT, in all Dicer immunoprecipitates (Fig. 5A). In addition, we cross-compared our data with a published Dicer proteomic screen and noted a strong overlap (54 hits are in the top 100 published hits), further confirming the robustness of our screen.(*59*) Principal component analysis showed distinct grouping of Dicer versus control immunoprecipitates (Fig. S5B). However, control versus ADAR1-depleted conditions did not group separately, suggesting that minimal changes in the Dicer interactome occurred during self-dsRNA sensing. Consistently, there were no significant changes in Dicer interactors after ADAR1 depletion, irrespective of ruxolitinib treatment (Fig. S5C-D). The four most enriched (yet non-significant) Dicer interactors after ADAR1 depletion were not enriched in ruxolitinib-treated cells, suggesting that the enrichment resulted from increased mRNA and protein abundance (Fig. S5E). Finally, GO enrichment analyses of the Dicer interactome between conditions showed no major changes (Fig. S5F). We conclude that Dicer’s role in self-dsRNA sensing does not involve major protein complex remodelling.

**Figure 5:**
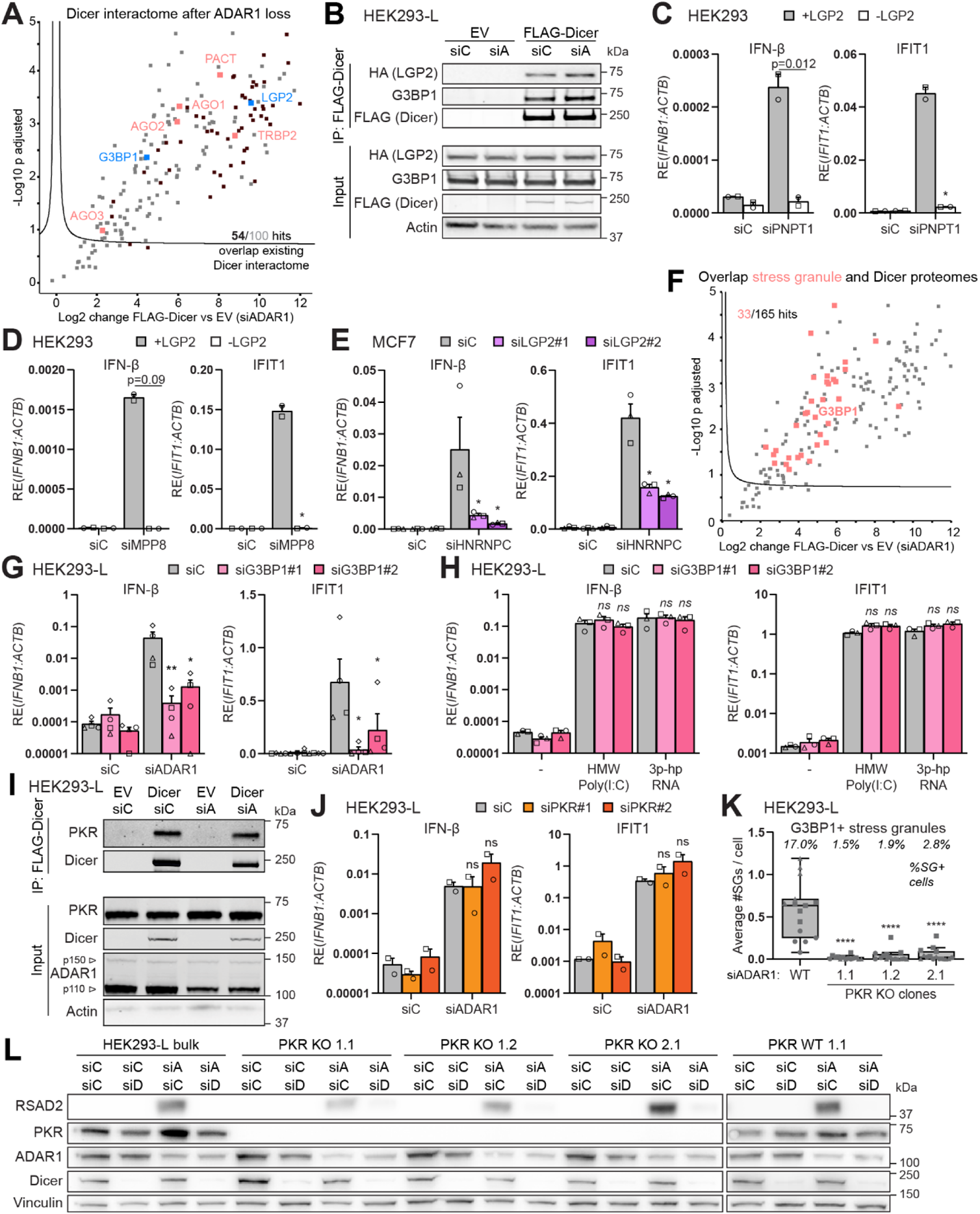
Dicer interactors LGP2 and G3BP1 are required for self-dsRNA sensing, independent of PKR and stress granules. **(A)** Volcano plot of Dicer interactome in HEK293-L cells 72h post-siADAR1 (n=3). Black datapoints indicate overlap with top-100 Dicer interactors of a published dataset.(*59*) Curved line indicates significance boundary. **(B)** Co-IP of FLAG-Dicer with selected interactors from (A), analysed by immunoblot. **(C-D)** HEK293 and HEK293-L cells analysed by RT-qPCR (N=2). **(C)** 4 days post-siPNPT1 transfection. **(D)** 6 days post-siMPP8 transfection. **(E)** RT-qPCR of MCF7 cells 72h post-siRNA transfection (n=3). **(F)** Volcano plot of Dicer interactome in HEK293-L cells 72h post-siADAR1, shown in Fig. 5A (n=3). Red datapoints indicate overlap with stress granule proteome.(*97*) **(G and J)** RT-qPCR of HEK293-L cells 72h post-siRNA transfection (n=3). **(H)** RT-qPCR of HEK293-L cells pretreated for 24h with siRNAs, then stimulated for 16h with HMW poly(I:C) or 3p-hpRNA (n=3). **(I)** Co-IP of FLAG-Dicer in siRNA-treated HEK293-L cells, analysed by immunoblot (n=3). **(K)** Quantification (15 images per condition) of stress granule formation shown in Fig. S7C of HEK293-L wt and *EIF2AK2* (encoding PKR) knockout clones 72h post-siADAR1 and IFN-I treatment (n=3). **(L)** HEK293-L wt and *EIF2AK2* (encoding PKR) knockout clones 72h post-siRNA transfection analysed by immunoblot (n=3). Data shown as means ±SEM (RT-qPCRs) or representative blots. Box plots showing interquartile range (IQR), median and whiskers (1.5×IQR), shapes represent biological repeats. Statistical tests: paired t-test (C-D), one-way ANOVA (E, G, H and and J-K). ns, not significant; \**P*<0.05; **P<0.01; ****P<0.0001. EV, empty vector; siA, siADAR1; siPN, siPNPT1; siC, siControl.

### Dicer interactor LGP2 is broadly required for the IFN-I response to self-dsRNA

Given the lack of overt changes in the Dicer interactome upon ADAR1 depletion, we hypothesized that Dicer must be engaged in homeostatic interactions with other proteins that are required for self-dsRNA sensing. Therefore, we examined the Dicer interactome for interactors contained within the GO term for regulators of the IFN-I response (Fig. S5G). The strongest hit was LGP2 (that is overexpressed in HEK293-L cells), which was previously identified as a Dicer interaction partner.(*40*) Co-immunoprecipitation (co-IP) of FLAG-Dicer and HA-LGP2 confirmed that Dicer associates with LGP2 at steady-state and in ADAR1-deficient cells (Fig. 5B and S6A). As LGP2 is essential for MDA5-mediated IFN-I induction in ADAR1-deficient cells, we examined whether LGP2, like Dicer, is required for self-dsRNA sensing in other contexts as well.(*5*) Indeed, the IFN-I response induced in HEK293 cells by mt-dsRNA and nuclear-derived self-dsRNA following depletion of PNPase or MPP8, respectively, required the presence of LGP2 (Fig. 5C-D and S6B-D). Similarly, knockdown of LGP2 in MCF7 cells attenuated the IFN-I response caused by hnRNPC depletion (Fig. 5E and S6E). Thus, Dicer interacts with LGP2 and both are broadly required for self-dsRNA sensing.

### Dicer and G3BP1 act on self-dsRNA sensing independent of PKR and stress granules

Interestingly, the SG-nucleator G3BP1 is also a Dicer interactor that is contained in the GO-term for regulators of the IFN-I response due to the putative role of SGs in the regulation of dsRNA sensing (Fig. S5G).(*50*) Co-IP confirmed the interaction between FLAG-Dicer and endogenous G3BP1 in control and ADAR1-depleted cells (Fig. 5B). Furthermore, the Dicer interactome shows a considerable overlap with the SG proteome, namely, 33 of 165 significant Dicer interactors (Fig. 5F). The dsRNA sensors LGP2, MDA5, RIG-I and PKR localize to SGs after infection by various RNA viruses, but whether SGs perform regulatory roles in the dsRNA response – and whether positive or negative – is a matter of ongoing debate and is likely context-dependent.(*50, 52, 60–65*) We therefore postulated that Dicer and its interactors could act on self-dsRNA sensing through SGs. While SG formation has been reported to occur in ADAR1-deficient cells, it is unclear whether they perform a regulatory role during self-dsRNA sensing.(*51, 52*) As expected, G3BP1 depletion attenuated SG induction in ADAR1-deficient A549-L cells (Fig. S6F), similar to our observations for Dicer depletion (Fig. 3B-C, 3E, 3F-G and S3B and E). Interestingly, co-depletion of G3BP1 attenuated the IFN-I response in ADAR1-deficient cells (Fig. 5G and S6G-H). Furthermore, depletion of G3BP1 did not affect IFN-I induction following transfection of exogenous RLR ligands HMW Poly(I:C) or 3p-hpRNA (Fig. 5H and S6I), in line with our observations for Dicer depletion (Fig. 2A).

To further investigate if Dicer and G3BP1 act on self-dsRNA sensing through SGs, we investigated the early events of SG nucleation. DsRNA-induced SG formation is initiated by PKR activation.(*50*) While PKR was not identified as an interactor of Dicer in our proteomic screen, others reported that Dicer binds and inhibits PKR during Sindbis virus infection.(*59*) Using adjusted lysis conditions, we recapitulated the Dicer-PKR interaction in co-IP experiments (Fig. 5I). To investigate how PKR impacts on the IFN-I response and SG formation, we co-depleted PKR and ADAR1 from HEK293-L cells. Surprisingly, PKR depletion using two-independent siRNAs did not abrogate the IFN-I response in cells lacking ADAR1 (Fig. 5J and S7A-B). To substantiate these findings, we made use of CRISPR/Cas9 to generate three individual *EIF2AK2* (encoding PKR) knockout clones using two independent PKR gRNAs. As expected, SG induction was lost in *EIF2AK2* knockout clones depleted of ADAR1 and treated with recombinant IFN-I (Fig. 6K and S7C). However, in line with the above findings, the IFN-I response upon ADAR1 depletion was not abrogated in all three knockout clones (Fig. 5L and S7D). Importantly, Dicer co-depletion still attenuates the IFN-I response upon ADAR1 ablation in *EIF2AK2* knockout cells. These results demonstrate that PKR and SGs are not required for self-dsRNA sensing, indicating that Dicer and G3BP1 act independently of the PKR-SG axis.

**Figure 6:**
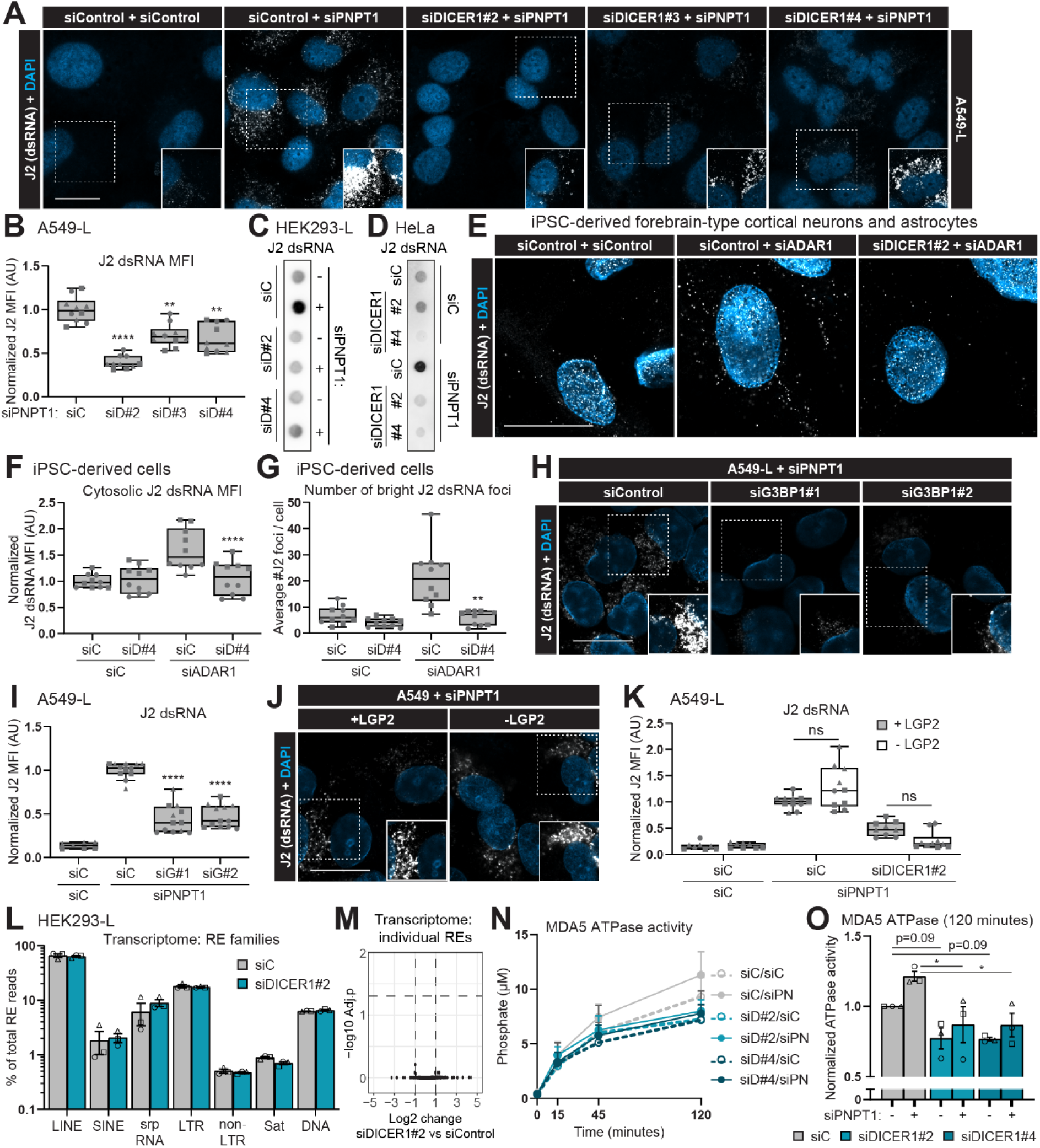
Dicer and G3BP1 regulate the accumulation of immunostimulatory self-dsRNA. **(A-B, H-K)** dsRNA accumulation in A549-L cells 72h post-siRNA transfection (n=3). **(A, H and J)** Visualized by J2-dsRNA immunofluorescence. Inset shows increased exposure of dashed box. **(B, I and K)** dsRNA quantified by mean fluorescence intensity (MFI) per image, normalized to siPNPT1/siC. **(C-D)** J2 dsRNA dot-immunoblot of total RNA from HEK293-L or HeLa cells 96h post-siRNA transfection (n=3). **(E-G)** Healthy hiPSC-derived forebrain-type cortical neurons and astrocytes (differentiated as in Fig. 1I) 5 days post-siRNA transfection (n=2). **(E)** dsRNA accumulation visualized by J2-dsRNA immunofluorescence on proteinase-K treated cells to expose dsRNA epitopes. **(F)** dsRNA quantified by cytosolic MFI. **(G)** Quantification of J2+ foci with an intensity over 5 a.u. in Fig. S11A-B. **(L-M)** Repetitive element analysis of transcriptomics from HEK293-L cells 72h post-siRNA transfection (n=3). **(L)** Family-level classification of differentially expressed repetitive elements. **(M)** Volcano plot showing individual repetitive elements. **(N-O)** *In vitro* MDA5 activity after incubation with RNA extracted from HEK293-L cells 72h post-siRNA transfection, measured by ATP hydrolysis using malachite green to measure inorganic phosphate (n=3). **(N)** Time-course shown as means ±SEM. **(O)** Total ATPase activity quantified after 120 minutes. Data shown as means ±SEM (RT-qPCRs) or representative images/blots. Scale bars: 20 µm. Box plots showing interquartile range (IQR), median and whiskers (1.5×IQR), shapes represent biological repeats. Statistical tests: one-way ANOVA (B, M and O), paired t-test (F-G), two-way ANOVA (K). ns, not significant; \**P*<0.05; **P<0.01; \*\*\*\**P*<0.0001. siD, siDICER1; siPN, siPNPT1, siG, siG3BP1; siC, siControl.

### Dicer and G3BP1 are required for the accumulation of self-dsRNA

Since Dicer is required for MDA5/LGP2-driven IFN-I induction, PKR-mediated SG induction and OAS1-3/RNAse L driven RNA degradation in ADAR1-deficient cells, we hypothesized that Dicer most likely acts upstream in the self-dsRNA response. To determine whether Dicer acts directly on self-dsRNA accumulation, we used the J2 antibody that specifically recognizes dsRNA, with a strong preference for perfectly base-paired dsRNA of at least 40 nt in length.(*66, 67*) We visualized the accumulation of mt-dsRNA following PNPase depletion through immunofluorescence, revealing formation of J2+ foci around mitochondria (Fig. S8A), as described before.(*10, 13, 68*) Strikingly, Dicer co-depletion using three independent siRNAs significantly reduces mt-dsRNA accumulation after PNPase depletion in A549-L cells (Fig. 6A-B). Specifically, Dicer co-depletion reduces the intensity of J2+ foci in PNPase-deficient cells, while the total number of foci remained unchanged (Fig. S8B-C). Loss of mt-dsRNA accumulation after Dicer/PNPase co-depletion was recapitulated with immunofluorescence of HEK293-L cells by simultaneous CRISPR/Cas9-mediated knockout of *PNPT1* and *DICER1* (Fig. S8D). Consistently, dot blot analysis of total RNA isolated from siRNA-treated HEK293-L and Hela cells revealed a buildup of J2-reactive dsRNA in PNPase-depleted cells, but not in PNPase/Dicer co-depleted cells (Fig. 6C-D). In line with the observation above, we observed a buildup of cytosolic J2+ dsRNA foci after ADAR1 depletion in hiPSC-derived mixed neuron-astrocyte cultures, which was dependent on Dicer expression (Fig. 6E-G and S9A-C). Note that this required careful titration of proteinase K treatment to expose dsRNA epitopes while avoiding over-digestion (Fig. S9D). We attempted to measure Dicer’s impact on self-dsRNA buildup following ADAR1 or hnRNPC depletion in immortalized cell lines but we were unable to detect increased J2+ foci in these cells, regardless of proteinase K treatment (Fig. S8E-F). This discrepancy may reflect the abundant expression of long self-dsRNAs in neurons through alternative splicing, which has been linked to the high prevalence of neuroinflammatory symptoms in AGS patients.(*69*) Overall, we conclude that Dicer is required for the accumulation of self-dsRNA of both nuclear and mitochondrial origin.

Next, we investigated whether Dicer interactors G3PB1 and LGP2 act together with Dicer to control self-dsRNA accumulation. Indeed, G3BP1 co-depletion significantly reduces J2 dsRNA accumulation after PNPase depletion in A549-L cells, akin Dicer (Fig. 6H-I). In contrast, the presence or absence of LGP2 does not affect J2 dsRNA levels after PNPase depletion (Fig. 6J-K). We conclude that Dicer and G3BP1 are jointly required for self-dsRNA accumulation in cytoplasmic foci.

### Dicer regulates immunostimulatory self-dsRNA accumulation

We next tested whether Dicer’s catalytic activity is required for self-dsRNA accumulation by re-expressing wild-type (wt) or catalytically inactive Dicer in Dicer-depleted cells. HEK293-L cells stably expressing C-terminally V5-tagged, siRNA-resistant Dicer were generated using the PiggyBac system (siResDicer wt). Notably, siResDicer wt expression did not restore ISG induction after ADAR1 depletion or J2 dsRNA accumulation following PNPase depletion in siDICER1-treated cells, despite robust expression (Fig. S10A-B). These data suggest that immunostimulatory self-dsRNA accumulation may require a specific Dicer isoform or that ectopic Dicer expression does not fully mimic endogenous Dicer. As an alternative to looking directly at dsRNA processing, we first examined self-dsRNA fate using our transcriptome analysis of ADAR1-depleted HEK293-L cells (Fig. 1C). Nuclear-derived self-dsRNAs, like those accumulating upon loss of ADAR1, hnRNPC or MPP8, largely originate from REs.(*15–17, 22, 30, 70*) We noted that RE transcript levels were unaltered after Dicer depletion, both at the level of RE families and when looking at single RE transcripts (Fig. 6L-M). Unchanged RE transcript levels, coupled with reduced J2 dsRNA staining after Dicer depletion, suggests that Dicer influences the conformation or accessibility of the double-stranded structure of RE-derived RNAs, rather than their expression. We next assessed whether Dicer affects the immunostimulatory potential of self-dsRNAs using an *in vitro* MDA5 ATPase assay. As expected, RNA from PNPase-depleted cells modestly increased recombinant MDA5 activity (consistent with the weak IFN-I response observed after PNPase depletion) (Fig. 6N, O). In contrast, RNA from cells depleted of both Dicer and PNPase failed to increase MDA5 ATPase activity, indicating that loss of Dicer abrogated the accumulation of immunostimulatory self-dsRNA. Furthermore, RNA from cells depleted of Dicer alone resulted in a reduction in MDA5 ATPase activity, as compared to RNA from control HEK293-L cells. Altogether, we conclude that Dicer is required for self-dsRNA accumulation and its immunostimulatory capacity as a DAMP in inflammation.

## Discussion

We uncover a previously unrecognized role for human Dicer as a key driver of cell-intrinsic immunity to self-dsRNAs. Dicer enables the buildup of self-dsRNAs arising from nuclear and mitochondrial sources, which in turn engage dsRNA sensors such as the RLRs, PKR, and OAS1-3. Acting independently of microRNA processing, Dicer interacts with G3BP1 to facilitate immunostimulatory self-dsRNA accumulation, and with LGP2 and PKR to enable self-dsRNA recognition. These findings elucidate the upstream mechanisms that govern self-dsRNA accumulation and sensing, revealing Dicer as a central regulator of self-dsRNA homeostasis. This has broad implications for autoinflammatory diseases, cancer and infection where self-dsRNAs act as danger signals.

The evolutionary conservation of this mechanism is noteworthy. In *Drosophila,* innate immune responses to *Adar* ablation is similarly rescued by *Dicer-2* depletion, despite differences in downstream signalling pathways.(*71*) This conservation suggests that Dicer homologues have maintained the capacity to recognize self-dsRNA across species, though the extent of its role in other organisms remains to be determined.

Our findings implicate Dicer as a necessary factor for the generation of immunostimulatory self-dsRNAs. This is consistent with earlier findings that >80% of Dicer-bound RNAs are not pre-miRNAs, and include RE-derived and mitochondrial transcripts (the proposed self-dsRNA sources in our models), which are processed inefficiently by Dicer. (*17, 30, 42*) While we do not observe impact of Dicer on RE-transcript abundance, we do observe decreased levels of dsRNA in Dicer-depleted cells, suggesting that Dicer directly impacts on the formation or stabilization of RNA in an immunostimulatory, double-stranded conformation.

We propose three non-mutually exclusive models to explain Dicer’s involvement: (1) Dicer may stabilize self-dsRNA in its double-stranded conformation, (2) Dicer may function as a nucleic acid annealer, hybridizing (partially) complementary RNA strands to form self-dsRNA, and (3) Dicer may process imperfect self-dsRNAs to make double stranded regions more accessible and immunostimulatory.

The intramolecular secondary structure formed by RE, such as hairpin loops, are imperfect ligands for dsRNA sensors.(*3*) *In vitro*, Dicer’s DUF283 domain catalyzes dsRNA formation from complementary RNAs, including those with complex secondary structures that otherwise hinder base-pairing. (*72–74*) Dicer may remodel the secondary structure of REs to enable annealing of complementary elements into bona fide dsRNAs. In addition, Dicer may process imperfect self-dsRNAs into more immunostimulatory forms. This is supported by the inefficient processing of long dsRNA observed for full length human Dicer, which instead preferentially processes hairpin loops and imperfectly base paired dsRNA, like pre-miRNAs.(*36, 42, 59, 75–79*) While some studies observed processing of RE Alu transcripts into endogenous siRNAs by Dicer, careful interpretation of the data shows that reported processing efficiencies are low.(*55, 80, 81*) As such, Dicer may selectively recognize and remove hairpin loops and imperfectly base-paired regions in self-dsRNAs, thereby enriching more perfectly paired bona fide dsRNAs that serve as superior ligands for dsRNA sensors. Finally, Dicer may directly transfer bound self-dsRNAs to its interactors LGP2 and PKR, similar to miRNA transfer to Ago2.(*31*) Consistently, LGP2 binding to Dicer was shown to suppress Dicer degradation of dsRNA(*40*), supporting a model wherein Dicer stabilizes, anneals or processes self-dsRNAs, without degrading them.

While we observed no role for Dicer in sensing synthetic dsRNA ligands or viral RNA isolated from Dicer-proficient cells, this does not eliminate the possibility that Dicer contributes to dsRNA sensing during bona fide viral infections. An antiviral role for Dicer is supported by the fact that many viruses encode Dicer antagonists.(*34*) Future studies should carefully consider viral Dicer antagonists, potential limited RNAi and the possibility that Dicer-dependent self-dsRNAs function as damage-associated molecular patterns (DAMPs) during infection.

The requirement for G3BP1 but not SGs in the IFN-I response upon ADAR1 ablation highlights the importance of distinguishing between SG-dependent and independent functions of G3BP1. Although G3BP1 is the most abundant SG protein, only 18% of total G3BP1 localizes to these structures during stress.(*82*) Our finding that G3BP1 and Dicer interact and are jointly required for self-dsRNA accumulation suggests that G3BP1 may modulate Dicer substrate specificity as a co-factor. Consistently, known Dicer binding partners extensively modulate Dicer substrate specificity.(*83*)

The implications of dysregulated self-dsRNA sensing, and Dicers role therein, extends far beyond type I interferonopathies. Cytosolic mt-dsRNA is implicated in osteoarthritis, renal injury, Huntington’s disease, and cellular senescence, suggesting it functions as a bona fide DAMP.(*11–14*) Furthermore, RLR or PKR activation and elevated self-dsRNAs and IFN-Is are observed in amyotrophic lateral sclerosis, Alzheimer’s disease, type-I diabetes and systemic lupus erythematosus.(*14, 84–90*) In cancer, self-dsRNA-driven immunity enhances tumour susceptibility to immunotherapy.(*9, 91–96*)

The ability of Dicer depletion to rescue neuroinflammation in a hiPSC-derived AGS model, highlights therapeutic potential in type I interferonopathies and other disorders driven by aberrant self-dsRNA sensing. Our results suggest that modulating Dicer function could control pathological self-dsRNA responses without compromising antiviral immunity. Although direct targeting of Dicer is challenging due to its essential role in miRNA biogenesis, our discovery of co-factors that cooperate with Dicer in self-dsRNA homeostasis provides the option to target these co-factors or their interaction interface with Dicer.

In conclusion, our findings position Dicer as a central regulator of self-dsRNA homeostasis, beyond its canonical role in miRNA processing. The involvement of dysregulated self-dsRNA sensing in diverse pathological contexts – including neurodegeneration, autoimmunity, and cancer – suggests that these findings have broad implications for understanding and treating these diseases.

## Limitations of the study

Visualizing self-dsRNA accumulation posed technical challenges across cellular models. While J2 antibody staining confirmed Dicer-dependent dsRNA accumulation following PNPase depletion in multiple cell lines using siRNA and CRISPR-Cas9, dsRNA detection after ADAR1 loss was limited to hiPSC-derived neurons. The selective detection in neurons likely reflects higher basal levels of immunostimulatory dsRNAs, consistent with the prominent neurological manifestations of AGS. We were unable to visualize increased J2 signal following MPP8 or hnRNPC depletion, precluding direct assessment of Dicer’s impact on dsRNA accumulation in those contexts. Furthermore, rescue experiments using ectopic expression of a C-terminal tagged canonical Dicer isoform were unsuccessful, suggesting the involvement of an alternate isoform or a regulatory context that is not recapitulated by ectopic expression. Importantly, we validated our findings using three independent Dicer depletion strategies, thereby excluding the possibility of technique-specific off-target effects.

## Acknowledgements

We are grateful to Anna Hautekeete, Anna Swirk, and Shannah Gennesse for their assistance during their internships in the laboratory. We thank Dr. Caetano Reis e Sousa for generously providing recombinant MDA5, Karel van Leeuwen for providing PiggyBac plasmids, and Mariana Guimarães Ramos for her advice on hiPSC culture and differentiation. We would like to thank the LUMC microscopy facility for support with the microscopy experiments.

We thank Arnoud de Ru and Rayman Tjokrodirijo for their support with proteomics experiments. We appreciate all members of the Van der Veen group for their support, technical assistance, and critical discussions throughout the course of this study. We thank Dr. Pierre Maillard for critical reading of the manuscript.

## Author contributions

**L.R.H.** designed the study, performed experiments, analysed data, prepared figures, and wrote the manuscript. **S.L., A.C.K., and D.G.** designed and performed experiments and analysed data. **S.L.** performed proteomic analysis. **D.G.** performed transcriptomic analysis. **T.O.** contributed to experimental design and provided strategic insight. **J.W.** and **Y.A.** performed experiments. **M.W.P.** assisted with the quantification of microscopy images, **P.A.V.** assisted with the mass spectrometry, **W.M.C.R.M.** provided resources and support for hiPSC culture and differentiation, and **H.M.** performed and assisted with bioinformatic analysis. **A.G.V.** conceived and designed the study, supervised the project, acquired funding, and wrote the manuscript. All authors have read and edited the manuscript.

## Data and materials availability

All data supporting the findings of this study are available in the main text or the supplementary materials. Mass spectrometry proteomics data have been deposited to the ProteomeXchange Consortium via the PRIDE partner repository under the dataset identifier **PXD068566**. Transcriptomics data have been deposited to the European Genome-phenome Archive (EGA) under the dataset identifier **EGAD50000002044**. CellProfiler pipelines detailing the exact analysis routines for the different experiments have been deposited to Zenodo (**DOI: 10.5281/zenodo.17365482**). Plasmids, cell lines, and other unique materials generated in this study are available from the corresponding author upon reasonable request; some materials may require a materials transfer agreement (MTA).

## Declaration of generative AI in the writing process

During the preparation of this work the authors used Microsoft Copilot exclusively to revise existing manuscript texts. After using this tool, the authors reviewed and edited the content as needed and take full responsibility for the content of the published article.

## Supplementary Materials

### Materials and Methods

#### Cell culture and reagents

HEK293, A549, U2OS (gift from Alfred Vertegaal, LUMC), HeLa and MCF7 cells were cultured in Dulbecco’s modified Eagle’s medium (DMEM; Gibco). HT29 cells (gift from Jacques van Dongen, LUMC) were cultured in Roswell Park Memorial Institute (RPMI) 1640 medium. All culture media were supplemented with 9% heat-inactivated foetal bovine serum (FBS; Capricorn), 2 mM L-glutamine (Gibco) and 100 U/ml penicillin/streptomycin (Gibco). Cells were cultured at 37 °C and at 10% CO_2_ for cells cultured in DMEM, others at 5% CO_2_. Universal Type I IFN (IFN-I, 11200, PBL Assay Science) was used at specified concentrations. 3p-IVT-dsRNA was generated by in vitro transcription using the MEGAshortscript (AM1354, Ambion) of the positive control pTRI-RNA 18S template included in the kit, according to manufacturer’s instructions, yielding a 128 bp 3p-dsRNA. The RLR ligands high molecular weight (HMW) poly(I:C), and 3p-hpRNA were obtained from Invivogen. Total RNA extracted from EMCV-infected HeLa cells was prepared as described before.(*40*) RLR ligands were complexed with Lipofectamine 2000 (Thermo Fisher Scientific) at a 1:1 ratio according to the manufacturer’s instructions and used at the indicated dosages. Dipyridamole (D9766, Sigma-Aldrich) was added to the culture medium at 50 µm, 30 minutes prior to transfection with total RNA from EMCV-infected cells to suppress viral replication.(*107*) Sodium arsenite (S7400, Sigma-Aldrich) was dissolved in ddH_2_O and added at indicated dosages 1 hour prior to cell fixation.

#### Differentiation and siRNA transfection of hiPSC-derived neural progenitor cells

Human neural progenitor cells (NPCs) (gift from Willeke van Roon, LUMC) were generated as described before from a healthy control patient (hPSCreg LUMCi023-A).(*108*) NPCs were cultured in STEMdiff neural progenitor medium (STEMCELL Technologies) on culture plates coated for 45 minutes with Matrigel hESC-Qualified Matrix (Corning). Plates used for differentiation were coated overnight with Poly-D-lysine hydrobromide (PDLh, P6407, Sigma-Aldrich) at 4°C, followed by 2 hour coating at 37°C with Laminin (L2020, Sigma-Aldrich). Differentiation into forebrain-type cortical neurons and astrocytes was done by plating 7.5 × 10^5^ NPCs in a PDLh/laminin-coated 6-well plate well in STEMdiff neuron differentiation medium (STEMCELL Technologies), which was refreshed daily. After 1 week, 1.25 × 10^5^ cells were replated to PDLh/laminin-coated 24-well plates (containing coverslips for confocal microscopy experiments) in STEMdiff neuron differentiation medium, followed by replacing half of the medium with fresh BrainPhys Neuronal medium every 2 to 3 days for 12 days of maturation. BrainPhys Neuronal medium was supplemented with SM1, N2, BDNF, GDNF (#05795, STEMCELL technologies), 0.5 mg/ml Dibutyryl-cAMP (#73886, STEMCELL technologies), 35 ng/ml Ascorbic Acid (A4544, Sigma-Aldrich) and 100 U/ml penicillin/streptomycin (Gibco, Thermo Fisher Scientific). On maturation day 10, cells were transfected with 2 µl of each indicated siGenome siRNA (5 µM, Dharmacon) using a 1:1 ratio of siRNA to Lipofectamine RNAiMAX transfection reagent (ThermoFisher) and according to manufacturer’s instructions. Complete BrainPhys Neuronal Medium was replenished 4 hours after transfection. Differentiated hiPSC-derived forebrain-type cortical neurons and astrocytes were harvested on maturation day 16. Correct differentiation was verified by expression of neuronal marker MAP2 and astrocyte marker GFAP on immunofluorescence confocal microscopy and by RT-qPCR.

#### Plasmid transfection, retroviral transduction and PiggyBac transposition

The production of retroviral particles and transduction of HEK293 cells to introduce the stable expression of 3FLAG-LGP2 (HEK293-L) has been performed and described previously.(*5*) To generate A549 3FLAG-LGP2 cells (A549-L), and HEK293 3HA-LGP2 cells (HEK293-L, used in co-IP experiments), A549 and HEK293 cells were transduced the same way as described previously(*5*), and selected using 0.5 µg/ml of puromycin (Sigma-Aldrich).

Plasmids were transfected using Lipofectamine 2000 (Invitrogen, Thermo Fisher Scientific), according to manufacturer’s protocol. pCAGGS-FLAG-Dicer was a gift from Phil Sharp, (Addgene #41584(*109*)). A lentiviral plasmid encoding C-terminally V5-tagged Dicer was ordered from Horizon (CCSB-Broad LentiORF: OHS6085-213575385) and Genscript introduced five synonymous base changes in the siRNA target regions of both siDICER1#2 (CGAA[G>A]GA[A>G]GA[G>A]GC[T>C]GA[C>T]TATGAA) and siDICER1#3 (CACAGA[A>G]TA[T>C][C>A]GATC[C>G]TA[T>C]GTTCAA) to generate siRNA resistant siResDicer-V5. We subcloned siResDicer-V5 into a plasmid encoding the PiggyBac transposon system (a gift from Karel van Leeuwen).

To generate HEK293-L cells stably expressing ectopic wildtype Dicer, we transfected cells with plasmids of the PiggyBac transposon system encoding siResDicer or EV coupled with an internal ribosomal entry site (IRES) to GFP. Cells were maintained in culture for two weeks in order to lose transient plasmid-derived GFP expression. Subsequently, cells stably expressing siResDicer or EV construct were sorted based on GFP expression into a bulk population to generate HEK293-L PiggyBac siResDicer wt and EV cell lines.

#### Gene silencing using siRNAs and PMOs

For gene silencing using siRNAs, individual siGenome siRNAs were purchased at Dharmacon targeting *DICER1* (D-003483-01 to −04, corresponding to siDICER1#1-4) *ADAR1* (D-008630-04), *DHX58* (D-010582-01 and −04, corresponding to siLGP2#1 and 2 respectively), *G3BP1* (D-012099-05 and −018 corresponding to siG3BP1#1 and 2 respectively) and *EIF2AK2* (D-003527-03 and −4, corresponding to siPKR#1 and 2 respectively), SMARTpool siGenome siRNAs against *MPHOSPH8* (M-021680-00), *HNRNPC* (M-011869-01), *PNPT1* (M-019454-00), *DGCR8* (M-015713-01), *DROSHA* (M-016996-02), and a nontargeting control (D-001210-03, corresponding to siControl). Cells were transfected with 25 pmol/ml of each siRNA using DharmaFECT 1 (Dharmacon) according to the manufacturer’s instructions, or using Lipofectamine RNAiMAX transfection reagent for hiPSC-derived cells as described above.

For gene silencing using phosphoroamidate morpholino oligomers (PMOs), splice-modifying PMOs to induce an exon-skip and frame shift were designed and ordered from Gene Tools LLC. PMO sequences are listed in Table 1. PMOs were delivered via endocytic uptake by replenishing cells with medium lacking penicillin/streptomycin and containing 4 µM Endo-Porter PEG (Gene Tool LLC) and 10 µM of each PMO, or by using PMOs conjugated to a cell-penetrating octa-guanidine dendrimer ‘vivo’ moiety (vPMOs) that were added at 10 µM to complete medium.

#### CRISPR-Cas9 gene editing

For CRISPR-Cas9-mediated gene editing, Alt-R Cas9-GFP nuclease, Alt-R electroporation enhancer, tracrRNA and custom crRNAs were ordered at Integrated DNA Technologies (IDT) and prepared before nucleofection into HEK293-L cells using 4D-Nucleofector System (Lonza) according to manufacturer’s instructions. crRNA sequences are listed in Table 1. For transient knockout experiments of *DICER1* and *PNPT1*, bulk cultures were maintained for indicated time periods. Knockout efficiencies were determined using the ICE (Inference of CRISPR Edits) tool of Synthego on Sanger sequencing reads of genomic PCR products using primers listed in Table 1. For generation of stable *EIF2AK2* (encoding PKR) knockout clones, bulk population was rested for one day post nucleofection before single cell sorting Cas9-GFP+ cells into 96-well plates. Clones were expanded over three weeks before screening for *EIF2AK2* knockouts by ICE-analysis of gene editing and immunoblotting for PKR protein as stated below. One clone that was complete wildtype but went through the selection process was maintained as wildtype (WT1.1) control, alongside the bulk wildtype population.

#### RT-qPCR

Total RNA was extracted using TRIzol or TRIzol LS reagent (Invitrogen) according to the manufacturer’s protocol. For cDNA synthesis, 200-500 ng RNA was treated with ezDNAse (Invitrogen) for 10 minutes and reverse transcribed using SuperScript IV VILO Master Mix (Invitrogen) according to the manufacturer’s instructions, and diluted 30x with H_2_O. Transcript levels were determined with 2 µl diluted cDNA and in technical duplicates using PowerUp or PowerTrack SYBR Green Master Mix (Applied Biosystems) on a QuantStudio 3 (10 µL reactions) or QuantStudio 6 (5 µL reactions) (Thermo Fisher Scientific). The primers used for RT-qPCR amplification are listed in Table 1. Relative expression (RE) levels were calculated using β-actin (*ACTB*) expression levels as an internal reference. Where indicated, RE values were normalized to the expression level of the indicated control condition.

#### Immunoblotting SDS-PAGE and dot blots

Cells were lysed in SDS lysis buffer (1% SDS, 150 mM NaCl, 50 mM Tris-HCl pH 7.4) or 0.5% NP-40 lysis buffer (0.5% IGEPAL CA-630 (Sigma-Aldrich), 50 mM of Tris-HCl pH 7.4, 150 mM of NaCl, 5mM MgCl_2_, 10% glycerol), supplemented with EDTA-free cOmplete mini protease inhibitor (Roche) and Genius nuclease (Santa Cruz) at manufacturer’s recommended concentrations. For immunoblotting of phosphorylated proteins, Phosphatase Inhibitor Cocktail V (Merck) was added at manufacturer’s recommended concentration to lysis buffers. Lysates in 0.5% NP-40 lysis buffer were cleared by centrifugation (15 min, 14,500 g at 4°C). Protein concentrations were equalized after BCA Protein Assay (Pierce, Thermo Fisher Scientific).

Samples were resolved on 4-15% Mini-PROTEAN TGX precast gels (Bio-Rad) or NuPAGE™ or Bolt™ Bis-Tris, 4 to 12%, 1.0 mm Protein Gels and transferred to Immobilon-P PVDF (Millipore) or nitrocellulose (Bio-Rad) membranes using the Trans-Blot Turbo Transfer System (Bio-Rad). Membranes were blocked using 5% non-fat dry milk (NFDM) in PBS-T (PBS, 0.1% Tween-20) and incubated with the indicated primary and secondary antibodies, diluted in 1% NFDM in PBS-T or Western BLoT Immuno Booster 1 or 2 solution (Takara Bio). Antibodies and dilutions are listed in Table 2. PVDF membranes were developed on a ChemiDoc MP Imaging System (Bio-Rad) using Luminata Crescendo, or Luminata Forte ECL substrate (both Millipore). Nitrocellulose membranes were developed on an Odyssey CLX Imaging System (LI-COR Biosciences). When required for reprobing, blots were stripped using Restore PLUS Western Blot Stripping Buffer (Thermo Scientific) according to the manufacturer’s instructions. For dot blot analysis of dsRNA content in cells, total RNA extracts were equalized and 2 µl dots containing 500-2000 ng RNA were dotted onto an Amersham Hybond-N+ nylon membrane (Cytiva). Membranes were left at least 30 minutes until dry and crosslinked with 240 mJ/cm^2^ UVC (254 nm) followed by blocking 1 hour in 5% NFDM in PBS-T and 1 hour in SuperBlock TBS Blocking Buffer (37536, Thermo Scientific). Membranes were incubated overnight in SuperBlock with 1:1000 J2 dsRNA antibody (Jena Bioscience), washed and incubated with secondary antibody in SuperBlock before developing immunoblot as described before.

#### Semi-denaturing detergent agarose gel electrophoresis (SDD-AGE)

To detect MDA5 oligomers, cells were lysed in SDD-AGE lysis buffer (0.5% IGEPAL CA-630 (Sigma-Aldrich), 50 mM of Tris pH 7.4, 150 mM of NaCl, 10% glycerol) supplemented with EDTA-free cOmplete mini protease inhibitor (Roche) for 30 min while rotating at 4°C. Lysates were cleared and protein concentrations were normalized as described above. Samples were incubated with 4× SDD-AGE sample buffer (2× TBE, 20% glycerol, 8% SDS, 0.01% bromophenol blue) at room temperature for 15 min and were resolved on vertical 1.5% agarose gels containing 1× TBE in 0.1% SDS in running buffer (1× TBE and 0.1% SDS), ran at 80 volts. Proteins were transferred onto Immobilon-P PVDF membranes (Millipore) and analysed by immunoblotting as described above.

#### FLAG immunoprecipitation

HEK293 3HA-LGP2 cells (HEK293-L) were transfected with control or ADAR1 siRNAs. Culture medium was refreshed after 8 hours, for indicated samples 1 mM Ruxolitinib (Invivogen) was added. 24 hours post-siRNA transfection, cells were transfected with pCAGGS-3FLAG-Dicer or empty vector (EV) plasmids. 72 hours post-siRNA transfection, cells were washed with ice cold PBS and lysed for 30 min at 4°C in NP-40 lysis buffer (50 mM Tris-HCl pH 7.5, 150 mM NaCl, 5 mM MgCl_2_, 10% ultrapure glycerol, 0.1% NP-40), supplemented with Complete-EDTA-free Protease Inhibitor Cocktail. Lysates were cleared by centrifugation for 15 minutes at 14.500 g at 4°C, normalized as described before, and then precleared for 1hr at 4°C with uncoupled Dynabeads protein G (Invitrogen). Dynabeads were blocked overnight in 1% BSA and monoclonal anti-FLAG M2 antibody (Sigma-Aldrich) was immobilized to blocked Dynabeads as per manufacturer’s instructions. Antibody-coupled beads were incubated with lysates for 4 hours at 4 °C. Beads were washed 3 times using lysis buffer and proteins were eluted using competitive inhibition with 0.5-1 mg/ml 1xFLAG peptide (synthesized in house by the LUMC Peptide Facility). Eluates were immunoblotted as described above, or used for downstream mass spectrometry analysis.

Co-IPs of PKR and Dicer were performed as described above, with the following alterations based on Montavon et al.(*59*) Cells were harvested in a Triton lysis buffer (50 mM Tris-HCl pH 7.5, 150 mM NaCl, 5 mM EDTA, 0.05% SDS, 1% triton) and treated for 20 minutes Turbo DNase (Invitrogen) at 37°C. Antibody-coupled beads were incubated with lysates overnight at 4 °C. Beads were washed 3 times using lysis buffer, 3 times using wash buffer (50 mM Tris-HCl pH 7.5, 200 mM NaCl, 5 mM EDTA, 0.05% SDS, 1% triton, supplemented with Complete-EDTA-free Protease Inhibitor Cocktail) and twice with ice cold PBS.

#### Transcriptomic analysis

Total RNA was extracted from HEK293-L cells 72 hours post-siRNA transfection, as described before. RNA libraries were prepared using the NEBNext Ultra II Directional RNA Library Prep Kit (New England Biolabs) following the manufacturer’s protocol (NEB #E7760S/L).

Ribosomal RNA depletion and library preparation was performed using the Qiagen fast select kit (#334376). Clustering and DNA sequencing using the NovaSeq6000 was performed according to manufacturer’s protocols. A concentration of 1.1 nM of DNA was used.

Raw sequencing reads were processed using the nf-core RNA-seq pipeline v3.18.0. In this pipeline, STAR (v2.7.11b)(*110*) was used to align reads to the GRCh38 human reference genome using settings --outFilterMultimapNmax 100 and --winAnchorMultimapNmax 100 to retain reads that map to multiple loci in the genome (in particular repetitive elements). The transcriptome index database was generated during pipeline execution using the combined Ensemble 113 and Dfam 3.7 TE annotation to define all genes. Salmon (v1.10.3)(*111*) was used for quantification of the mapped reads using libtype ISR and --gcBias to correct for GC biases.

Differential expression analysis was conducted using edgeR (v4.6.3) in R. Genes with a Benjamini and Hochberg FDR adjusted p-value < 0.05 and a log2 fold change > 1 were considered significantly differentially expressed.

GO enrichment analysis was performed on differentially expressed genes using clusterProfiler(v4.16.0) with the MSigDB Hallmark gene set. Enriched biological processes and pathways were identified using a false discovery rate (FDR) threshold of < 0.05.

Volcano plots and PCA plots were generated using ggplot2 in R. Heatmaps were generated using pheatmap in R. Transcriptomic data was visualized to highlight changes in coding genes and TEs across conditions. Family-level TE expression was visualized using ggplot2. Differential expression analyses data is available in Supplemental Table 1.

ISGs were defined based on a published list of 379 human ISGs from microarray data of multiple IFN treated and virus infected cell lines and tissues(*112*), which we further filtered by removing genes that were not enriched in transcriptomic analyses in a publication that treated human peripheral blood mononuclear cells with multiple type I interferons.(*113*) This resulted in a list of 252 ISGs, that is available in Supplemental Table 2.

The transcriptomics data have been deposited to the European Genome-phenome Archive (EGA) with the dataset identifier EGAD50000002044.

#### Proteomic analysis

For GeLCMS analysis, gel bands were reduced with 10 mM dithiothreitol, alkylated with 50 mM iodoacetamide. In-gel trypsin digestion was performed using a Proteineer DP digestion robot (Bruker). Tryptic peptides were extracted from the gel slices using 50/50/0.1 water/acetonitrile/formic acid (v/v/v), after which samples were lyophilized. The samples were dissolved in water/formic acid (100/0.1 v/v) and analyzed by on-line C18 nanoHPLC MS/MS with a system consisting of an Ultimate3000nano gradient HPLC system (Thermo, Bremen, Germany), and an Exploris480 mass spectrometer (Thermo). Peptides were injected onto a cartridge precolumn (300 μm × 5 mm, C18 PepMap, 5 um, 100 A, and eluted via a homemade analytical nano-HPLC column (30 cm × 75 μm; Reprosil-Pur C18-AQ 1.9 um, 120 A (Dr. Maisch, Ammerbuch, Germany). The gradient was run from 2% to 40% solvent B (20/80/0.1 water/acetonitrile/formic acid (FA) v/v) in 30 min. The nano-HPLC column was drawn to a tip of ∼10 μm and acted as the electrospray needle of the MS source. The mass spectrometer was operated in data-dependent Top 20 MS/MS mode, with a HCD collision energy at 30% and recording of the MS2 spectrum in the orbitrap, with a quadrupole isolation width of 1.2 Da. In the master scan (MS1) the resolution was 120,000, the scan range 400-1500, at standard AGC target @maximum fill time of 50 ms. A lock mass correction on the background ion m/z=445.12003 was used. Precursors were dynamically excluded after n=1 with an exclusion duration of 10 s, and with a precursor range of 20 ppm. Charge states 2-5 were included. For MS2 the first mass was set to 110 Da, and the MS2 scan resolution was 30,000 at an AGC target of 100% @maximum fill time of 60 ms.

Maxquant analysis was done with default settings using version 2.5.1.0. The database was the Homo sapiens minimal database (20.596 entries). Proteomic hits were filtered to identify confident Dicer interactors; only hits that had 6+ peptides detected in 2+ replicates of at least one condition, and had a peptide ratio over 1.5 in the Dicer vs EV pulldown were included. Protein quantification was performed using intensity-based absolute quantification (iBAQ). Proteomics data was visualized using Perseus. P-values were calculated in Perseus using T-test with 250 number of variations. The hyperbolic significance threshold was calculated using: -log₁₀(p-value) = c + s × log₁₀(fold change) (c = 0.6 and s = 0.1).

The mass spectrometry proteomics data have been deposited to the ProteomeXchange Consortium via the PRIDE partner repository with the dataset identifier PXD068566. Furthermore, the significant Dicer interactors are available in Supplemental Table 3.

#### In vitro MDA5 ATPase assay

Total RNA extracts from HEK293-L cells 96 hours post siRNA-mediated PNPase and or Dicer depletion were incubated at 37°C with 300 nM recombinant human MDA5 protein (produced in insect cells as described before (*40*)) and 1 mM ATP in assay buffer (20 mM HEPES pH 7.8, 100 mM NaCl, 1 mM glycerol, 2 mM MgCl_2_, 2 mM DTT, RNasin (Promega)). At indicated timepoints, samples were taken and quenched by mixing 1:1 with 20 mM EDTA in assay buffer. The concentration of inorganic phosphate released by MDA5 ATP hydrolysis was measured by absorbance at 620 nm of malachite green binding to phosphate ions using the Malachite Green Phosphate Assay Kit (Sigma-Aldrich), according to manufacturer’s instructions.

#### Bioanalyzer

To evaluate RNase L activation, total RNA integrity was assessed using the Agilent 2100 Bioanalyzer (Agilent Technologies) with the RNA 6000 Nano Kit, following the manufacturer’s protocol. RNase L activation was inferred from degradation of cellular RNA, visualized by the appearance of a smeared electropherogram profile. RNA integrity was quantified using RNA integrity number (RIN) to support visual observations.

#### Confocal microscopy

Cells were seeded in 8-well Millicell EZ slides (Millipore) and treated as described in the figure legends. Millicell slides used to culture HEK293 cells were coated for 10 minutes with Poly-L-Lysine hydrobromide to enhance cell adherence. hiPSC-derived cells were plated on PDLh-Laminin coated coverslips in 24-well culture plates, as described above. Cells were fixed and permeabilized for 10 minutes in methanol at −20°C. Whenever indicated, fixed cells were treated with 0.0005 (hiPSC-derived cells) or 0.0015 (other cells) U/ml Proteinase K Solution, RNA grade (25530049, Invitrogen) for 30 minutes at 37°C to exposed dsRNA epitopes covered by dsRNA-binding proteins. Fixed cells were stained with antibodies (listed in Table 2) diluted in TNB (TBS, 0.5% blocking reagent (#11096176001, Roche), 0.02% Thimerosal (Sigma-Aldrich)). Washing steps were performed using TBS-T (100 mM Tris-HCl pH 7.4, 150 mM NaCl, 0.05% Tween-20). Slides were embedded in DAPI-containing Prolong Gold antifade mountant (Invitrogen). Images were acquired on a Zeiss LSM900 confocal microscope at the LUMC Light and Electron Microscopy Facility. Representative images were acquired using the Airyscan super resolution detector with a 40x/1.3 Oil or 63x/1.4 Oil DIC objective and processed using Zeiss Zen software. All other images intended for quantification were acquired using a 10x/0.45 or 20x/0.8 objective. Nuclei were outlined by hand based on DAPI staining.

#### Microscopy image analysis

Original microscopy data was converted to single-image tiff files using an ImageJ macro. Subsequently images were analysed using CellProfiler v 4.2.8 (installer with Stardist included installed from: https://github.com/glencoesoftware/CellProfiler/releases/v4.2.80002-ai/). Nuclei were detected using RunStardist in CellProfiler, after resizing the images for optimal nucei segmentation performance. Small objects, not resembling nuclei were filtered out using a fixed size threshold.

SG quantification was performed using the EnhanceOrSuppressFeatures module to suppress the background and the ‘global robust background’ threshold was applied to identify the spots. To determine the number of spots per cell, the detected nuclei objects were expanded with a 200 pixel radius, or until they touched each other. Subsequently the expanded nucleus signal was used to relate the SGs to individual cells. SG+ cells were defined as cells that had at least 2 detected SGs to reduce false-positive cell numbers from mis-identified SGs.

Given a higher cytoplasmic G3BP1 staining in HEK293 cells we made use of an adaptive threshold of 100 pixel window size with the robust background threshold. Furthermore, to correct for variation in the intensity of the staining between replicates the robust background threshold was adjusted, but kept the same per replicate (namely 4 times the standard deviations for the first replicate and 5 SDs for the other replicates).

SGs in hiPSC-derived mixed neurons and astrocytes were manually quantified based on G3BP1 staining. For each condition and biological replicate, six images were acquired at 10× magnification and visually inspected to determine the number of SG-positive cells. Manual quantification was performed due to limitations in automated detection accuracy in these cell types.

To quantify the amount of J2 dsRNA signal in A549 cells both J2 dsRNA spot intensity and number were measured. To improve the detection of J2 dsRNA spots, the EnhanceOrSuppressFeatures function was used, subsequently spots were detected using a global Otsu threshold, with the same parameters for all images. The number of J2 spots per cell was determined as described above for the SG. Furthermore the J2 mean fluorescence intensity (MFI) of the full images and the mean intensity per individual cell area was determined. In order to normalize J2 MFI between repeat experiments, J2 MFI was normalized within repeats to the PNPase depleted sample.

hiPSC-derived neurons and astrocytes showed high J2 dsRNA signal in nuclei. To focus on dsRNAs that can activate cytosolic dsRNA sensors, only J2 sRNA spots in the cytoplasm were counted. After detection of the nuclei with Stardist, the signal was expanded and subsequently the nucleus was excluded from the object. For hiPSC-derived cells, J2 MFI was normalized within repeats to the siControl treated sample. Cellprofiler pipelines detailing the exact analysis routines for the different experiments have been deposited to Zenodo (DOI: 10.5281/zenodo.17365482).

#### Data and statistical analysis

All experiment replicate numbers refer to biological replicates, unless explicitly stated otherwise. Statistical analysis was performed using GraphPad Prism software (version 10.2.3). In case of RT-qPCR data, non-normalized data was log-transformed prior to statistical testing. Single comparisons were performed using a paired Student’s *t*-test. Multiple group analysis was performed using repeated measures one-way or two-way ANOVA, followed by Holm-Šidak’s *post hoc* test. Statistical significance is represented by: ns, not significant; \**P* value < 0.05; \*\**P* value < 0.01; \*\*\**P* value < 0.001; \*\*\*\**P* value < 0.0001.

---

**Supplemental Figure 1: belonging to Figure 1.**
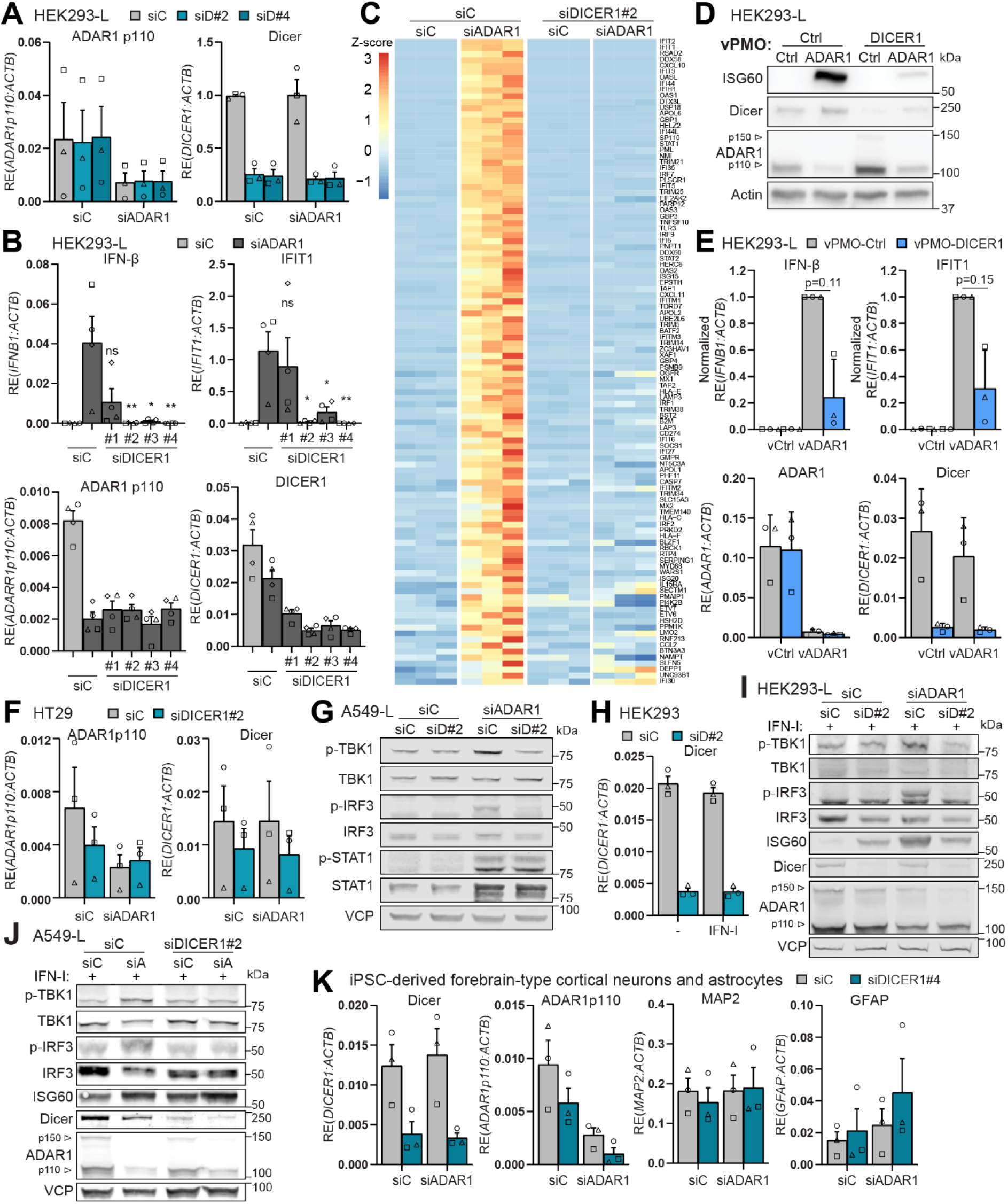
**(A)** RT-qPCR validation of ADAR1 and Dicer knockdowns in HEK293-L cells from Fig. 1A (n=3). ADAR1 knockdown verified using qPCR primers specific to ADAR1 p110 isoform, to circumvent upregulation of ADAR1 p150 isoform as an ISG, which would prevent assessment of knockdown efficiency. **(B)** RT-qPCR analysis of IFN-I response and knockdown efficiencies in ADAR1/Dicer-depleted HEK293-L cells from Fig. 1B (n=3). **(C)** Heatmap of all significant ISGs in siADAR1 vs siControl comparison, by transcriptomics of Fig. 1C (n=3). **(D-E)** HEK293-L cells treated with vPMOs targeting ADAR1 and Dicer (n=3). **(D)** Protein levels by immunoblot. **(E)** Transcript levels by RT-qPCR using qPCR primers that specifically amplify the exons targeted by the splice-modifying vPMOs **(F)** RT-qPCR validation of ADAR1 and Dicer knockdown in HT29 cells from Fig. 1E (n=3). **(G)** Phospho-immunoblot of A549-L cells shown in Fig. 3A, 72h post-transfection with siRNAs targeting ADAR1 and Dicer (n=3). **(H)** RT-qPCR validation of Dicer knockdown in HEK293-L cells from Fig. 1G (n=3). **(I-J)** Phospho-immunoblot of HEK293-L and A549-L cells 72h post-siRNA transfection, and recombinant IFN-I treatment to upregulate dsRNA sensors (n=3). **(K) h**iPSC-derived forebrain-type cortical neurons (MAP2) and astrocytes (GFAP) of Fig. 1I, harvested 5 days post-siRNA transfection. Differentiation and siRNA-knockdown efficiency verified by RT-qPCR. Data shown as means ±SEM (RT-qPCRs) or representative images/blots. Statistical tests: one-way ANOVA (B), paired t-test (E). \**P* < 0.05; \*\**P* < 0.01. siD, siDICER1; siC, siControl.

**Supplemental Figure 2: belonging to Figure 2.**
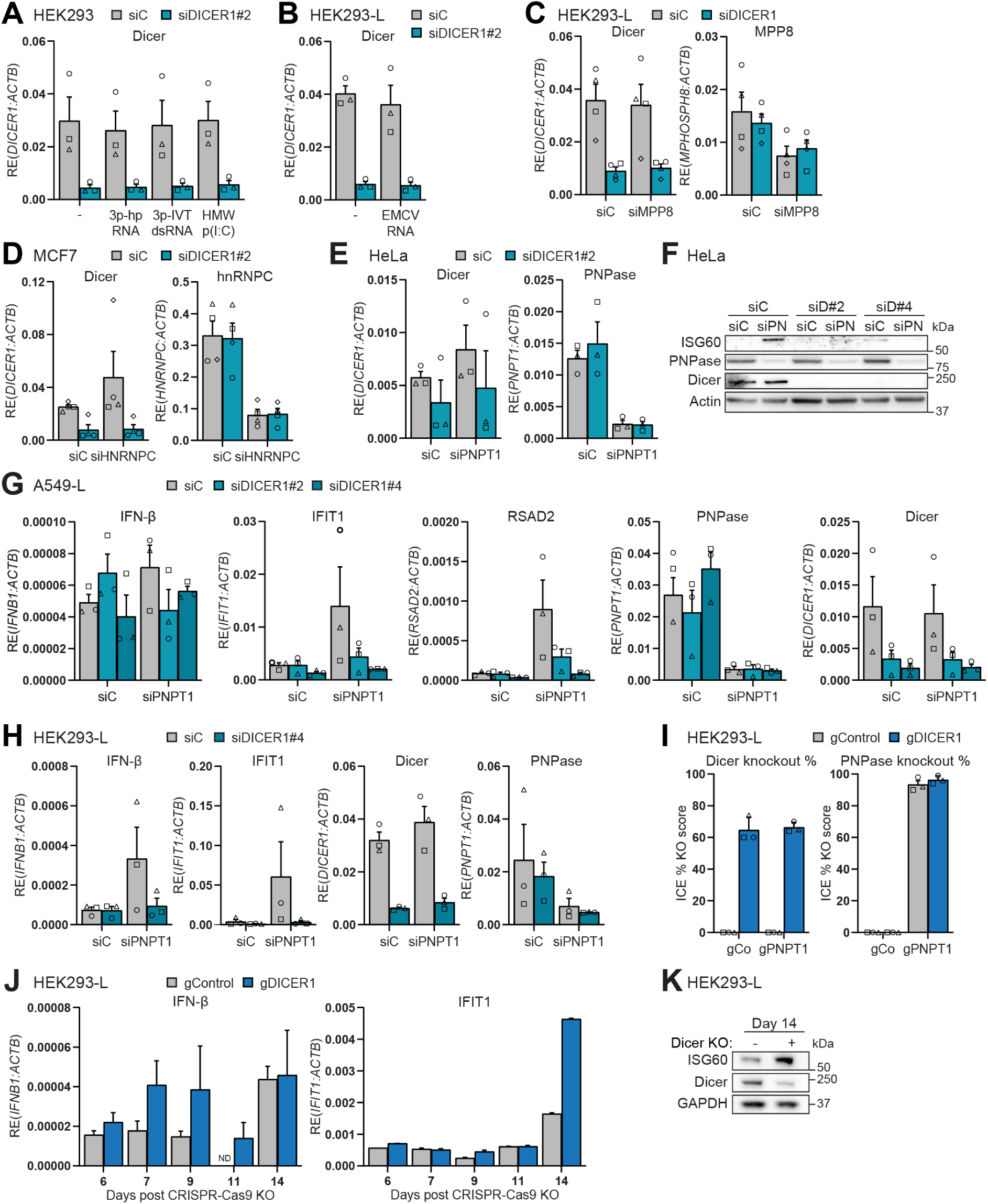
**(A-E)** RT-qPCR validation of siRNA knockdowns in HEK293 cells (**A**, n=3, related to Fig. 2A), HEK293-L cells (**B**, n=3, related to Fig. 2B; **C**, n=4, related to Fig. 2C), MCF7 cells (**D**, n=4, related to Fig. 2E) and HeLa cells (**E**, n=3, related to Fig. 2G). **(F)** HeLa cells 96h post-siRNA transfection analysed by immunoblot (n=1). **(G)** RT-qPCR analysis of A549-L cells shown in Fig. 2H, 96h post-siRNA transfection (n=3). **(H)** HEK293-L cells 96h post-siRNA transfection targeting PNPase and Dicer (n=3). **(I)** CRISPR-knockout efficiency percentage assessed by ICE (Inference of CRISPR Edits) analysis of samples shown in Fig. 2I-J (n=3). **(J-K)** HEK293-L cells 6-14 days post-nucleofection with Cas9 and gRNAs targeting Dicer (n=1). **(J)** IFN-I response by RT-qPCR. **(K)** Protein levels by immunoblot. ND, not detected. Data shown as means ±SD of technical duplicates (J) or as means ±SEM of biological repeats (other panels) (RT-qPCRs) or representative images/blots.

**Supplemental Figure 3: belonging to Figure 3.**
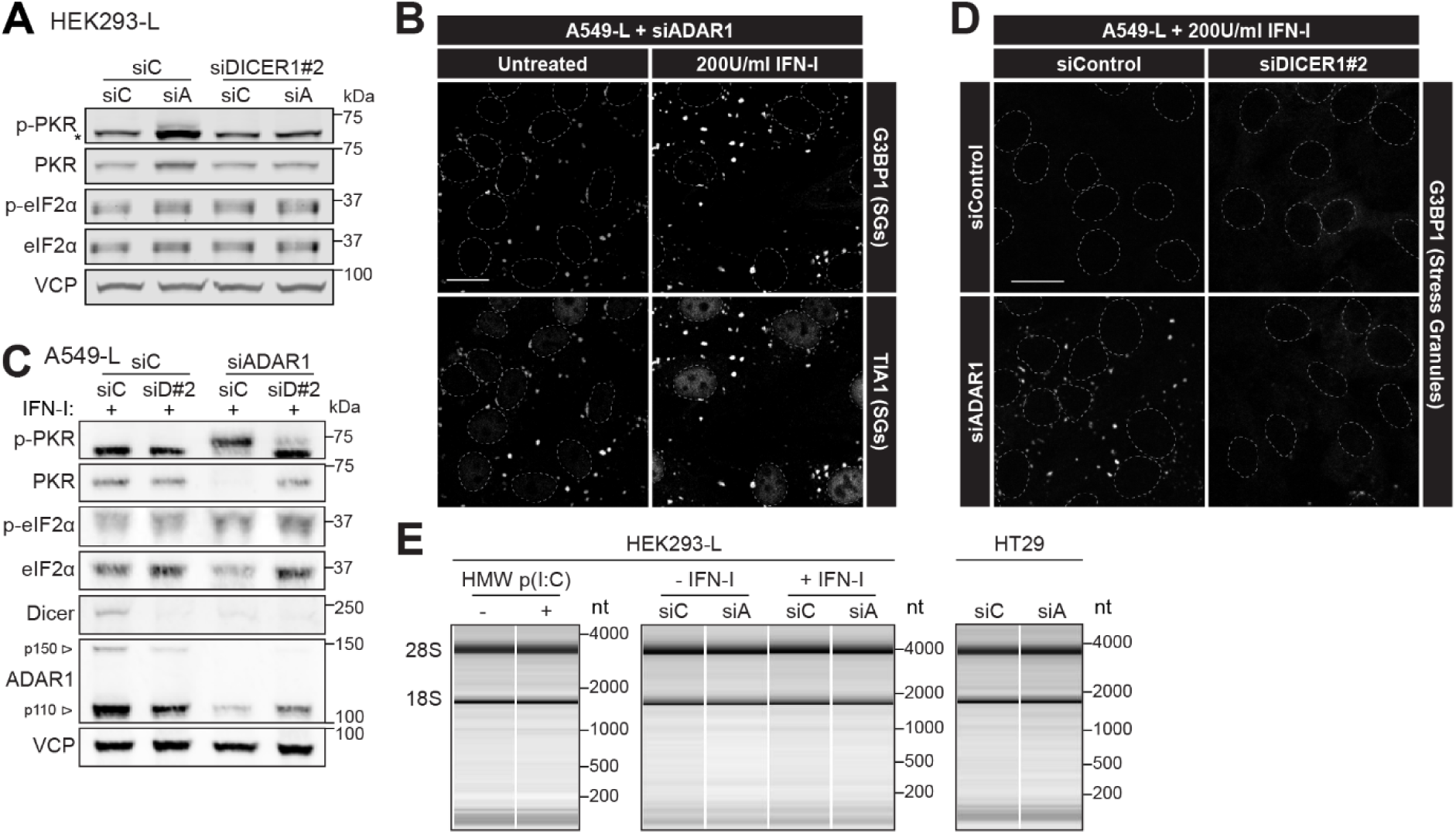
**(A)** Phospho-immunoblot of HEK293-L cells shown in Fig. 1F, 72h post-transfection with siRNAs targeting ADAR1 and Dicer (n=3). **(B)** Immunofluorescence of siRNA-treated A549-L cells stained after 72h for G3BP1 and the stress granule-exclusive marker TIA1. (n=2). **(C)** Phospho-immunoblot of A549-L cells, 72h post-siRNA transfection and IFN-I treatment to upregulate PKR (n=3). **(D)** Immunofluorescence of G3BP1 to visualize stress granules in A549-L cells 72h post-siRNA transfection and 48h post-recombinant IFN-I treatment to boost PKR expression (n=3). **(E)** HEK293-L and HT29 cells, 72h post-siRNA or 6h post-HMW poly(I:C) transfection. HEK293-L cells were treated with or without IFN-I 24h post-siRNA. OAS1-3/RNase L activation measured by integrity analysis of total RNA extracts using Bioanalyzer electrophoresis (n=2). Data shown as representative images/blots. Nuclear DAPI stains outlined, scale bars: 20 µm. siA, siADAR1.

**Supplemental Figure 4: belonging to Figure 4.**
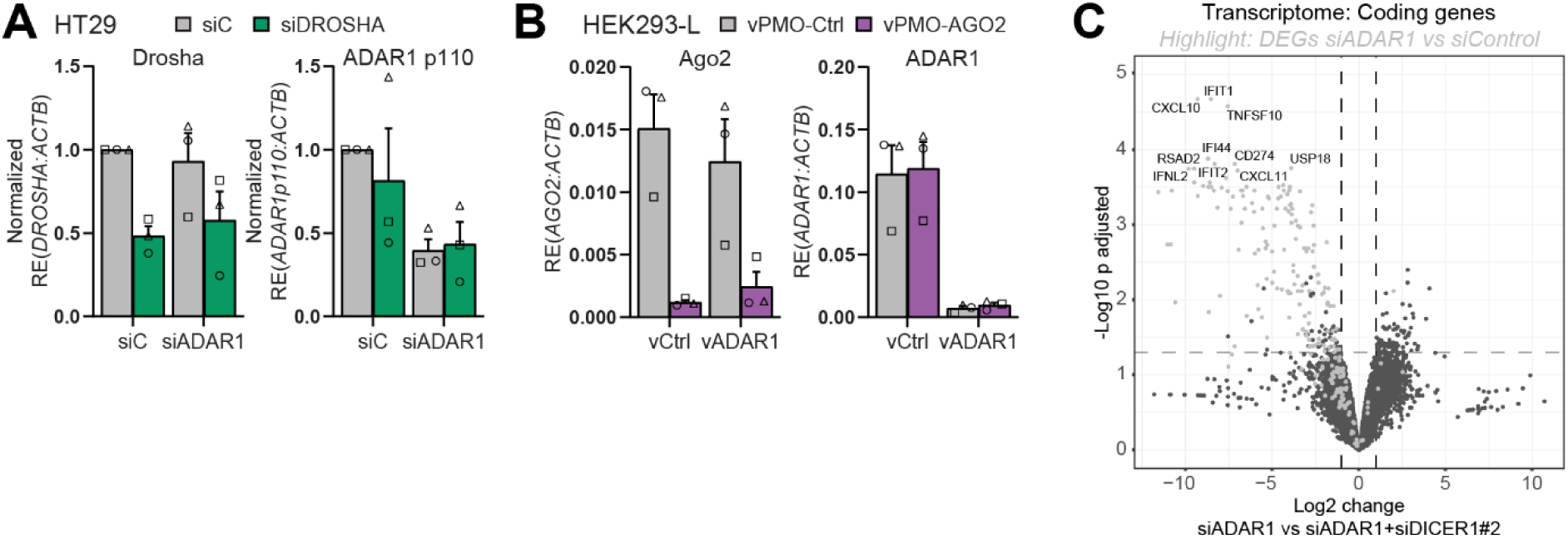
**(A)** RT-qPCR (means ±SEM) validation of siRNA-knockdown in HT29 cells from Fig. 4B (n=3). **(B)** RT-qPCR (means ±SEM) validation of vPMO-knockdown in HEK293-L cells from Fig. 4C, using qPCR primers that specifically amplify the exons targeted by the splice-modifying vPMOs (n=3). **(C)** Differential expression analysis of transcriptomics on HEK293-L cells shown in Fig. 1C, 72h post-siRNA transfection (n=3). Significantly differentially expressed genes (DEGs) in siADAR1 vs siControl comparison are highlighted in the volcano plot of the siADAR1 vs siADAR1+ siDICER1#2 comparison, to emphasize DEGs that are the result of the additional siDICER1#2 treatment.

**Supplemental Figure 5: belonging to Figure 5.**
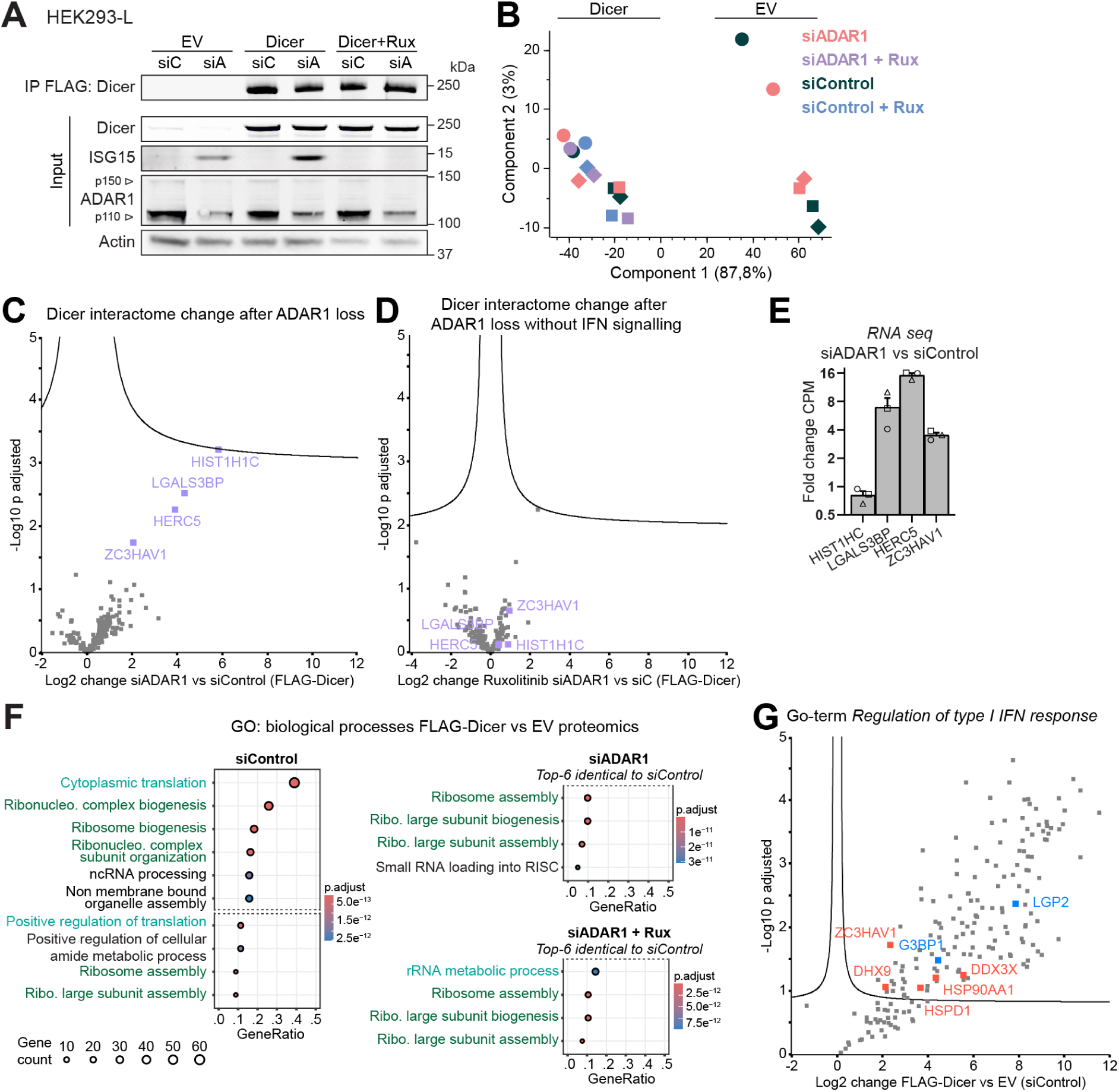
**(A)** Validation of FLAG-Dicer immunoprecipitation (used for proteomics) in HEK293-L cells treated with siControl, siADAR1, or siADAR1 + ruxolitinib (n=3). Input and IP fractions analysed by immunoblot. EV, empty vector; siA, siADAR1; Rux, ruxolitinib. **(B)** Principal component analysis of proteomics samples from (A); biological replicates indicated by matching symbols. **(C-D)** Volcano plots showing changes in Dicer interactome 72h after siADAR1 treatment (n=3). Enriched Dicer interactors in siADAR1 treated samples are highlighted in purple. **(C)** Enriched Dicer interactors after siADAR1 treatment, versus siControl. **(D)** Enriched Dicer interactors in siADAR1 + ruxolitinib treatment, versus siControl. **(E)** Transcript levels of enriched Dicer interactors after siADAR1 treatment in (C) (n=3). Transcript levels shown as the fold change of the counts per million (CPM) after siADAR1 treatment from transcriptomics shown in Fig. 1C of HEK293-L cells, 72h post-siRNA transfection. **(F)** GO-enrichment analysis of Dicer interactors 72h post-siRNA transfection. Color-coded by GO-term similarity; ribosome-related GO-terms are labelled green and translation-related GO-terms are labelled blue. **(G)** Volcano plot of Dicer interactome after 72h in siControl-treated HEK293-L cells (n=3). Interactors overlapping with the GO term *Regulation of type I IFN response* are named and highlighted red, or blue for those chosen for further study. Curved line in volcano plots indicates the significance boundary based on a hyperbolic threshold combining fold change and p-value criteria.

**Supplemental Figure 6: belonging to Figure 5.**
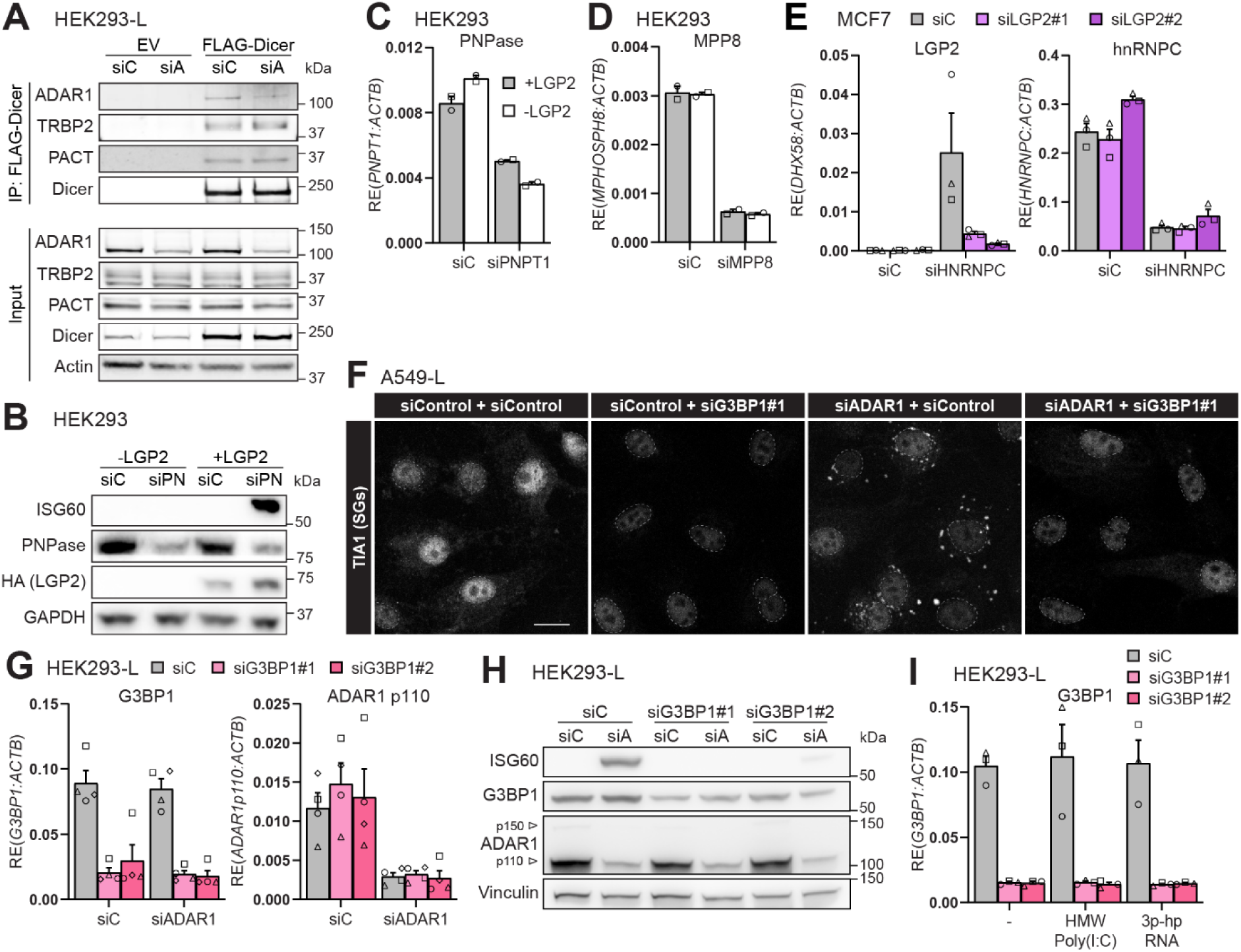
**(A)** Co-IP of FLAG-Dicer in siRNA-treated HEK293-L cells shown in Fig. 5B (n=3). Input and IP fractions analysed by immunoblot for known Dicer interactors. **(B-C)** siPNPT1-treated HEK293 and HEK293-L cells from Fig. 5C (N=2). **(B)** Protein levels analysed by immunoblot. **(C)** siRNA-knockdowns validated by RT-qPCR. **(D)** RT-qPCR validation of MPP8 knockdown in HEK293 and HEK293-L cells from Fig. 5D (N=2). **(F)** Immunofluorescence of A549-L cells stained for stress granule marker TIA1, 72h post-siRNA transfection (n=3). Scale bar: 20 µm. **(G)** RT-qPCR validation of siRNA-knockdowns in HEK293-L cells from Fig. 5G (n=3). **(H)** Immunoblot of HEK293-L cells shown in Fig. 5G, 72h post-siRNA transfection (n=1). **(I)** RT-qPCR validation of siRNA-knockdowns in HEK293-L cells from Fig. 5H (n=3). Data shown as means ±SEM (RT-qPCRs) or representative images/blots. EV, empty vector; siA, siADAR1; siC, siControl; siPN, siPNPT1.

**Supplemental Figure 7: belonging to Figure 5.**
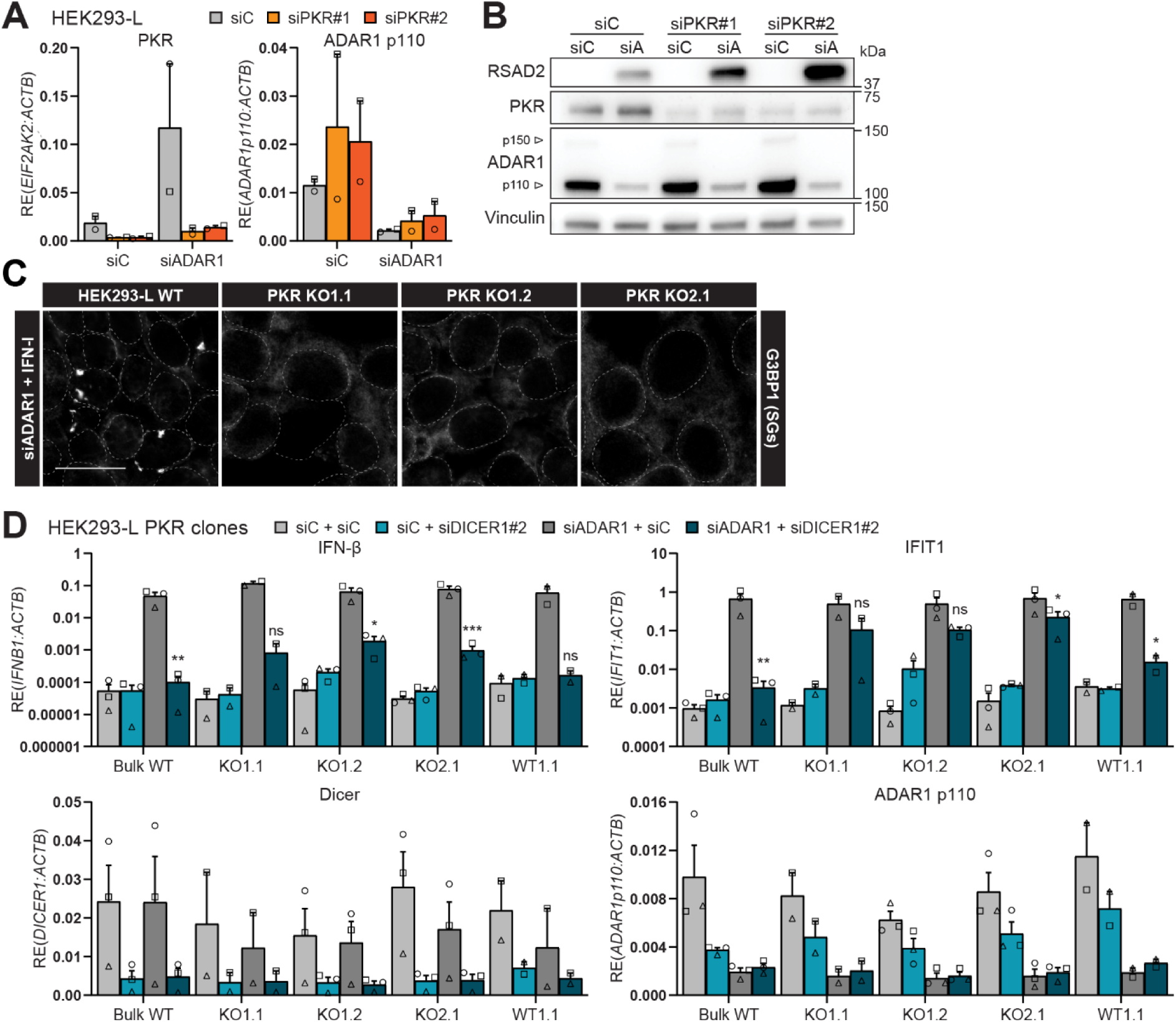
**(A-B)** HEK293-L cells shown in Fig. 6C, 72h post-siRNA transfection (n=2). **(A)** IFN-I response determined by RT-qPCR. **(B)** Protein levels assessed by immunoblot. **(C)** Stress granule formation in HEK293-L wt and *EIF2AK2* (encoding PKR) knockout clones 72h post-siADAR1 and IFN-I treatment (n=3). Visualized by G3BP1 immunofluorescence, quantified in Fig. 5K. Scale bar: 20 µm. **(D)** HEK293-L wt or *EIF2AK2* (encoding PKR) knockout clones shown in Fig. 5L, 72h post-siRNA transfection monitored by RT-qPCR (n=3, or n=2 for KO1.1 and WT1.1). Data shown as means ±SEM (RT-qPCRs) or representative images/blots. Statistical test: one-way ANOVA (D). ns, not significant; \**P*<0.05; **P<0.01; \*\*\**P*<0.001; \*\*\*\**P*<0.0001. siA, siADAR1; siD, siDICER1; siC, siControl.

**Supplemental Figure 8: belonging to Figure 6.**
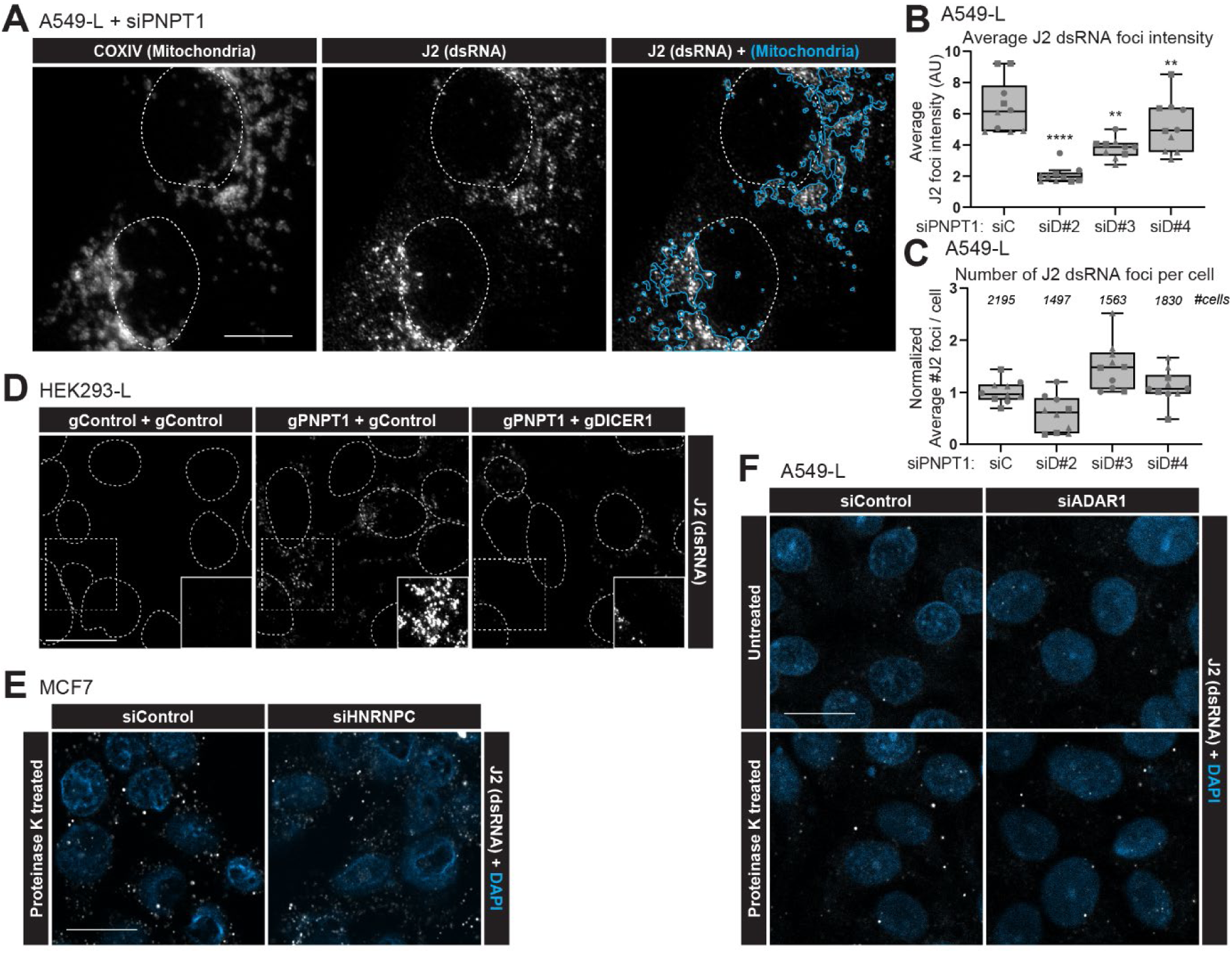
**(A)** Immunofluorescence of A549-L cells 72h post-siRNA transfection with siPNPT1 or siControl. Overlap of J2 dsRNA accumulation with mitochondria (outlined manually in blue)(n=2). Scale bar: 10 µm. **(B-C)** Quantification of J2 dsRNA foci intensity in **(B)** and number in **(C)** of A549-L cells shown in Fig. 7A, 72h post-siRNA transfection (n=3). **(D)** dsRNA accumulation in HEK293-L cells 5 days post-nucleofection with Cas9 and gRNAs targeting *PNPT1* and/or *DICER1*, visualized by J2-dsRNA immunofluorescence (n=1). Area outlined by dashed box shown overexposed in solid-line box. Scale bar: 20 µm. **(E-F)** J2 dsRNA immunofluorescence of siRNA-treated A549-L cells in **(E)** and MCF7 cells in **(F)** with or without proteinase K treatment to expose dsRNA epitopes covered by dsRNA-binding proteins (n=1). Scale bar: 20 µm. Data shown as representative images/blots. Box plots showing interquartile range (IQR), median and whiskers (1.5×IQR), shapes indicate biological repeats. Nuclear DAPI stains outlined (white).

**Supplemental Figure 9: belonging to Figure 6.**
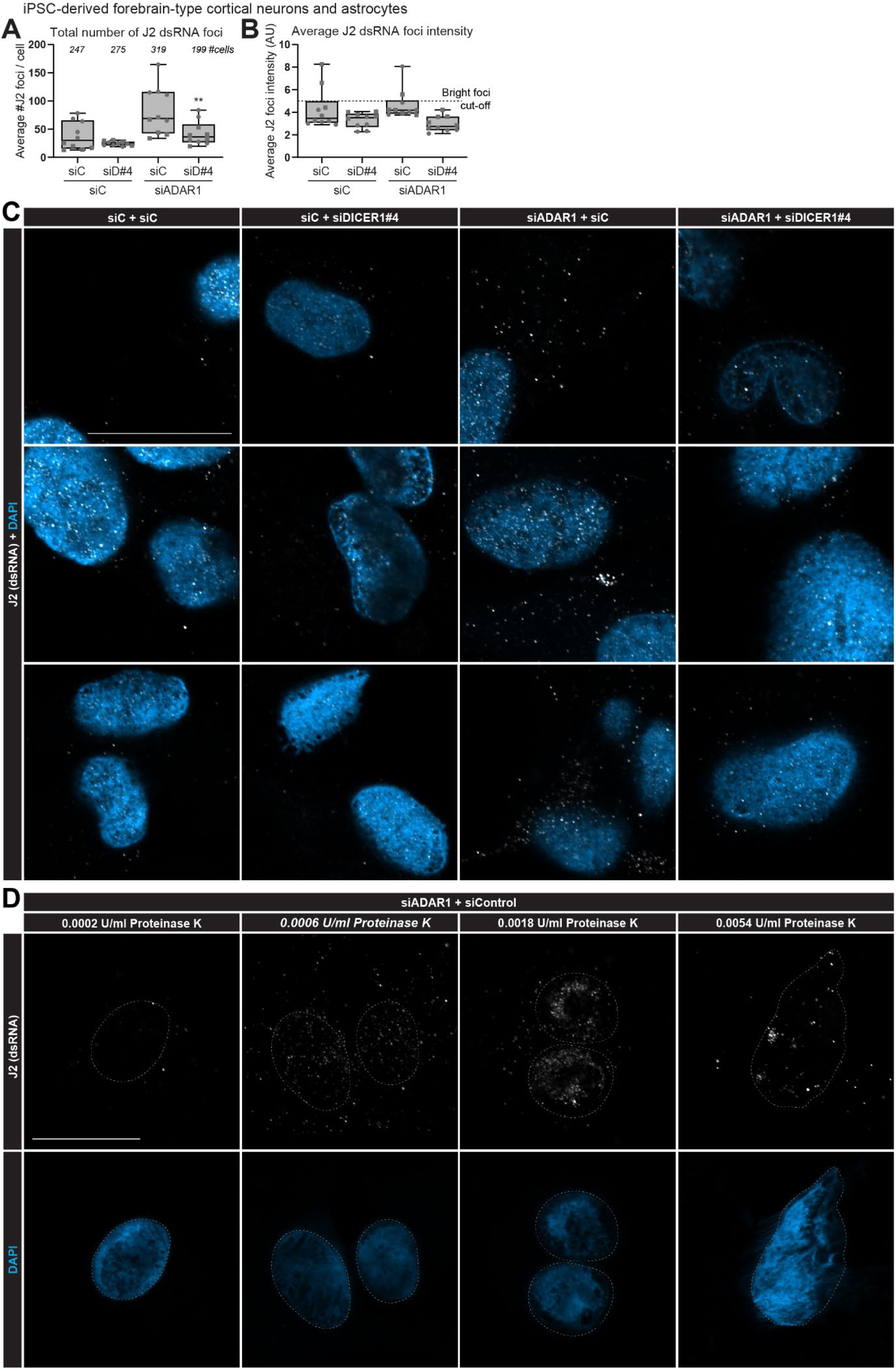
**(A-D)** hiPSC-derived forebrain-type cortical neurons (MAP2) and astrocytes (GFAP) of Fig. 7E, harvested 5 days post-siRNA transfection. dsRNA accumulation visualized by immunofluorescence using J2 antibody (n=2). **(A)** Quantification of total number of J2+ dsRNA foci. **(B)** Quantification of J2+ dsRNA foci average intensity (A.U.), dashed line shows cut-off for bright J2+ foci with an intensity over 5 a.u. that are displayed in Fig. 7G. **(C)** Additional representative images of dsRNA accumulation shown in Fig. 7E. **(D)** Proteinase K treatment was titered on hiPSC-derived cells to visualize cytoplasmic J2-dsRNA by exposing dsRNA epitopes otherwise covered by dsRNA-binding proteins, but without over digesting cells. 0.0006 U/ml was chosen for final experiments. Data shown as representative images. Box plots showing interquartile range (IQR), median and whiskers (1.5×IQR), shapes indicate biological repeats. Scale bars: 20 µm.

**Supplemental Figure 10: belonging to Figure 6.**
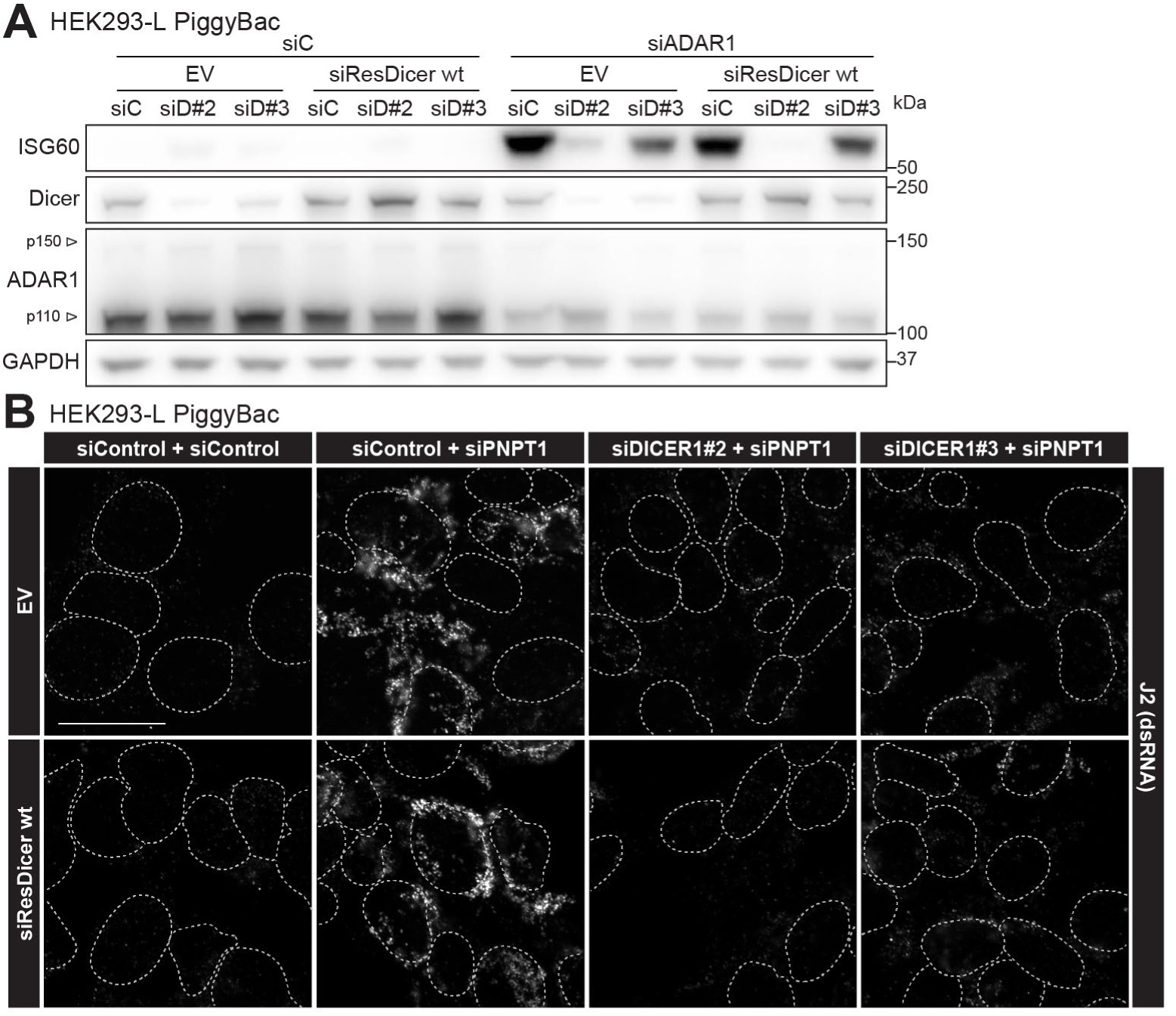
**(A-B)** HEK293-L cells stably expressing EV or siRNA-resistant siResDicer wt, 72h post-siRNA transfection (n=1). **(A)** Protein levels assessed by immunoblot. **(B)** J2 dsRNA accumulation assessed by immunofluorescence. Data shown as representative images. Nuclear DAPI stains outlined, scale bars: 20 µm. siD, siDICER1.

**Table S1:**
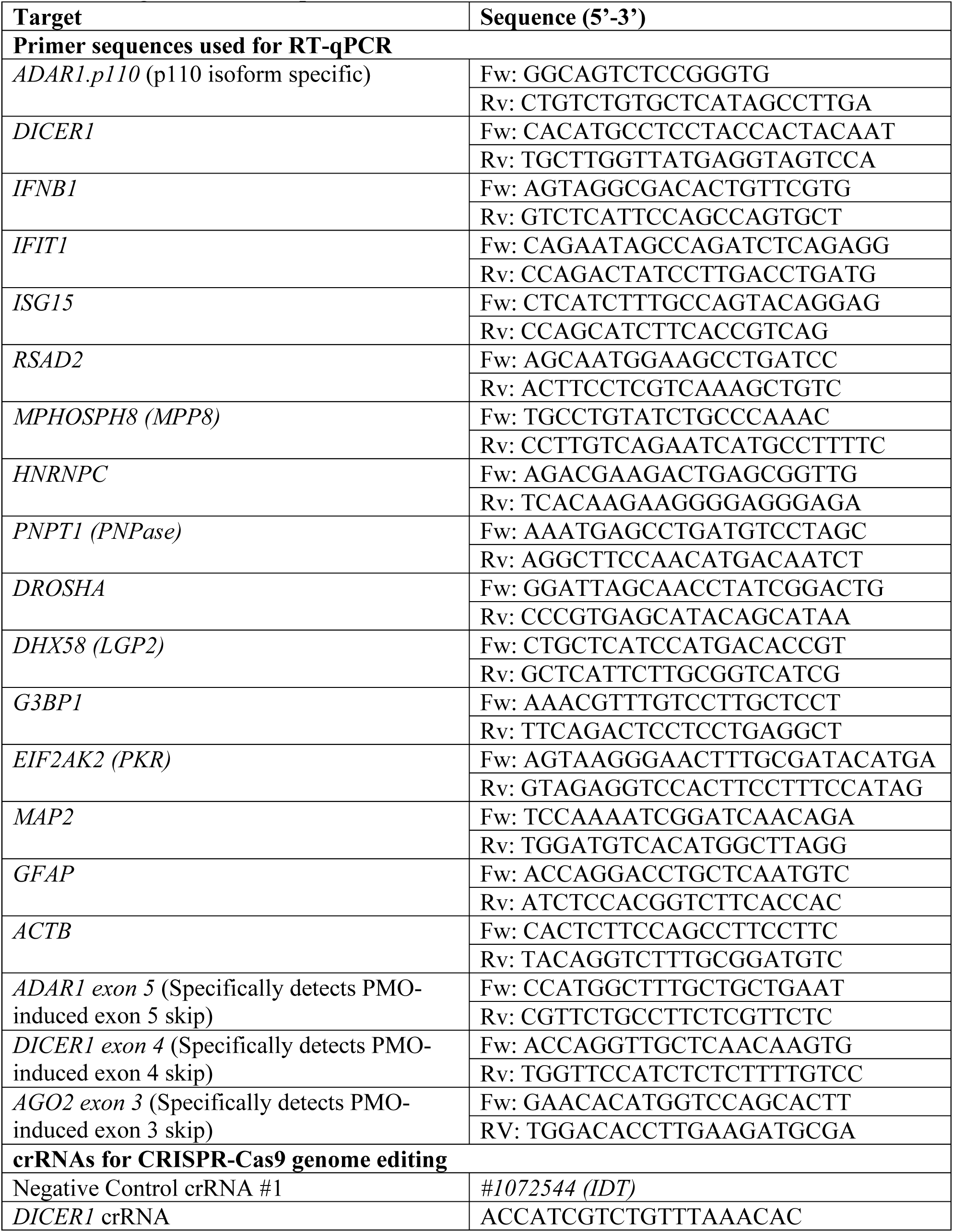
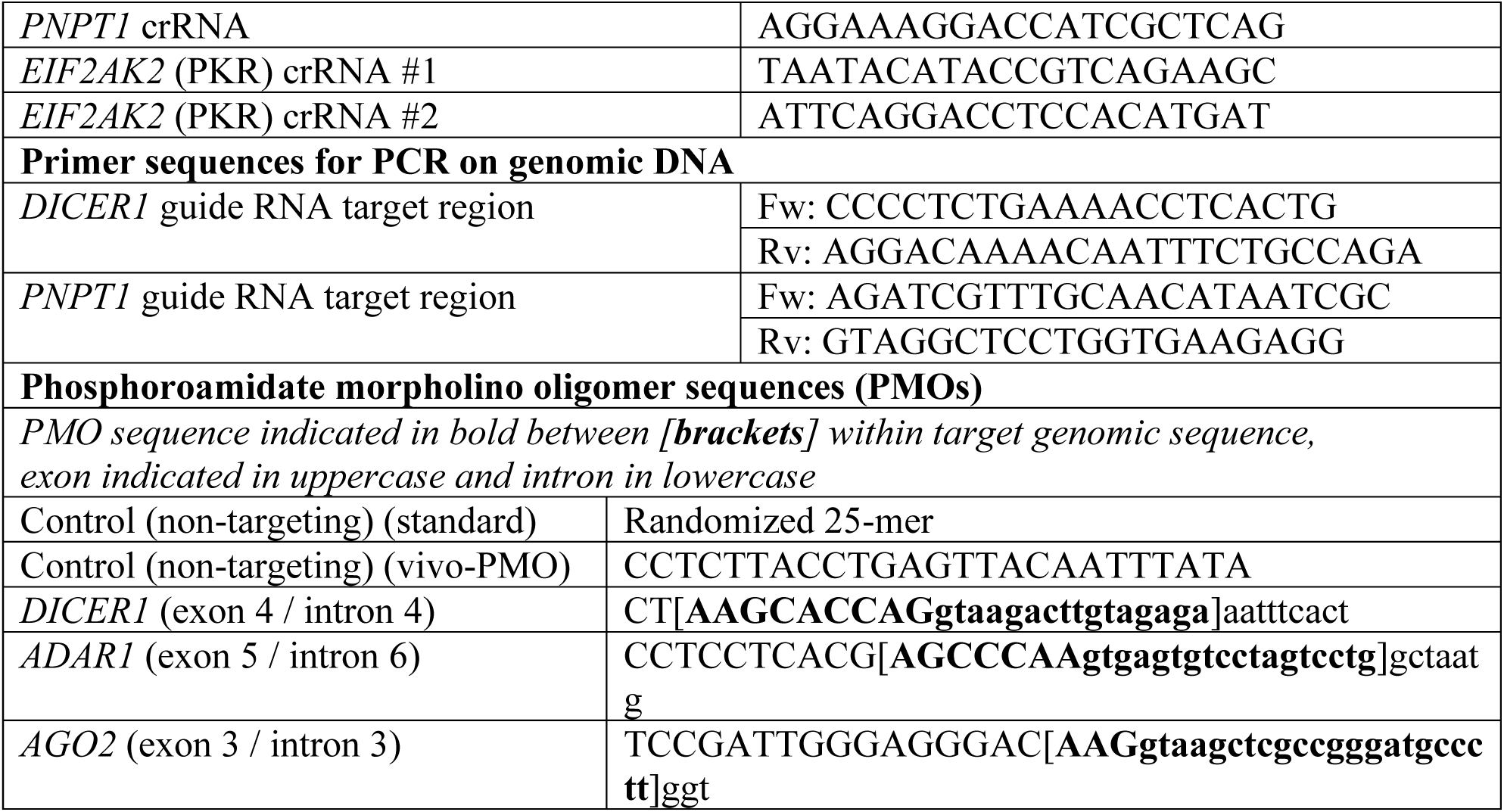
Oligonucleotide sequences.

**Table S2:** Antibodies.

| Target/Epitope | Application | Dilution | Catalog no. and company |
|---|---|---|---|
| DICER1 | WB | 1:1000 | 5362 (Cell Signaling Technology) |
| DICER1 | WB | 1:2000 | 68375-1-Ig (ProteinTech) |
| ADAR1 | WB | 1:500 | sc-73408 (Santa Cruz) |
| ISG60 (IFIT3) | WB | 1:500 | sc-393512 (Santa Cruz) |
| RSAD2 | WB | 1:1000 | 13996S (Cell Signaling Technology) |
| ISG15 | WB | 1:500 | sc-166712 (Santa Cruz) |
| MDA5 | WB | 1:1000 | 21775-1-AP (ProteinTech) |
| Phospho-TBK1 (Ser172) | WB | 1:500 | 5483 (Cell Signaling Technology) |
| TBK1 | WB | 1:1000 | 3504 (Cell Signaling Technology) |
| Phospho-IRF3 (Ser396) | WB | 1:500 | 4947 (Cell Signaling Technology) |
| IRF3 | WB | 1:1000 | 4302 (Cell Signaling Technology) |
| Phospho-STAT1 (Tyr701) | WB | 1:1000 | 7649 (Cell Signaling Technology) |
| STAT1 | WB | 1:1000 | 9172 (Cell Signaling Technology) |
| Phospho-PKR (Thr446) | WB | 1:1000 | ab32036 (Abcam) |
| PKR | WB | 1:1000 | 3072 (Cell Signaling Technology) |
| Phospho-eIF2α (Ser51) | WB | 1:1000 | 9721 (Cell Signaling Technology) |
| eIF2α | WB | 1:1000 | sc-133227 (Santa Cruz) |
| IκBα | WB | 1:1000 | 4814 (Cell Signaling Technology) |
| MPP8 | WB | 1:20,000 | 16796-1-AP (ProteinTech) |
| hnRNPC | WB | 1:2500 | sc-32308 (Santa Cruz) |
| PNPase | WB | 1:10,000 | 14487-1-AP (ProteinTech) |
| Drosha | WB | 1:1000 | 3364 (Cell Signaling Technology) |
| DGCR8 | WB | 1:1000 | 10996-1-AP (ProteinTech) |
| Ago2 | WB | 1:1000 | ab57113 (Abcam) |
| G3BP1 | WB/IF | 1:500/100 | sc-365338 (Santa Cruz) |
| G3BP1 | WB/IF | 1:1000 | 13057-2-AP (ProteinTech) |
| TIA1 | IF | 1:200 | 610352 (BD biosciences) |
| TRBP2 | WB | 1:500 | sc-514124 (Santa Cruz) |
| PACT | WB | 1:2500 | ab75749 (Abcam) |
| MAP2 | IF | 1:5000 | ab5392 (Abcam) |
| GFAP | IF | 1:5000 | 13-0300 (Invitrogen) |
| COXIV | IF | 1:250 | 11967 (Cell Signaling Technology) |
| FLAG | IP | 10 µg/IP | F1804 (Sigma-Aldrich) |
| J2 (dsRNA) | Dot/IF | 1:1000/200 | RNT-SCI-10010200 (Jena Bioscience) |
| VCP | WB | 1:5000 | 60316-1-Ig (ProteinTech) |
| GAPDH-HRP | WB | 1:10.000 | HRP-60004 (ProteinTech) |
| β-actin-HRP | WB | 1:5000 | sc-47778 HRP (Santa Cruz) |
| vinculin-HRP | WB | 1:5000 | sc-73614 HRP (Santa Cruz) |
| FLAG-HRP | WB | 1:20,000 | A8592 (Sigma-Aldrich) |
| HA-HRP | WB | 1:3,000 | 12013819001 (Roche) |
| FLAG-IRDye 800CW | WB | 1:10,000 | 600-431-383 (Rockland) |
| Goat-anti-mouse IgG-HRP | WB | 1:5000 | G-21040 (Invitrogen) |
| Goat-anti-rabbit IgG-HRP | WB | 1:5000 | 4050-05 (Southern Biotech) |
| Goat-anti-mouse IgG IRDye 680LT | WB | 1:5000 | 926-68020 (LI-COR Biosciences) |
| Goat-anti-rabbit IgG IRDye 800CW | WB | 1:5000 | 926-32211 (LI-COR Biosciences) |
| Goat-anti-rabbit Alexa488 | IF | 1:1000 | A-11034 (Invitrogen) |
| Goat-anti-rabbit Alexa568 | IF | 1:1000 | A-11036 (Invitrogen) |
| Goat-anti-rabbit Alexa647 | IF | 1:1000 | A-21245 (Invitrogen) |
| Goat-anti-mouse-IgG2a Alexa488 | IF | 1:1000 | A-21131 (Invitrogen) |
| Goat-anti-mouse-IgG2a AlexaPlus555 | IF | 1:1000 | SA5-10370-AFP555 (Invitrogen) |
| Goat-anti-mouse-IgG1 Alexa488 | IF | 1:1000 | A-21121 (Invitrogen) |
| Goat-anti-mouse-IgG1 Alexa568 | IF | 1:1000 | A-21124 (Invitrogen) |
| Goat-anti-mouse-IgG1 Alexa647 | IF | 1:1000 | A-21240 (Invitrogen) |
| Donkey-anti-rat-IgG AlexaPlus488 | IF | 1:1000 | A48269TR (Invitrogen) |
| Goat-anti-chicken-IgY AlexaPlus647 | IF | 1:1000 | A32933 (Invitrogen) |

*WB: Western Blot, Dot: Dot blot, IF: Immunofluorescence, IP: Immunoprecipitation*

**Data S1. (separate file)**

Supplemental Data 1 – Differential expression analyses transcriptomics

**Data S2. (separate file)**

Supplemental Data 2 - A list of 252 ISGs

**Data S3. (separate file)**

Supplemental Data 3 - Significant Dicer interactors proteomics

